# Mutually Repressive Transcription Factors Drive Intestinal Maturation for Weaning

**DOI:** 10.64898/2026.11.04.764083

**Authors:** M.K. Scales, R. Karsan, J. Underwood, A.M. Bara, Y. Wang, K. Sumigray, T. Lechler

**Affiliations:** Department of Cell Biology, Duke University, Durham, NC 27710; Department of Genetics, Yale University, New Haven, CT; Department of Dermatology, Duke University, Durham, NC 27710

## Abstract

In mammals, rapid neonatal growth depends on specialized intestinal enterocytes that absorb nutrients from milk. These cells exhibit distinct functions, morphologies, and gene expression programs that support neonatal nutrition before transitioning to an adult state at weaning. However, the transcriptional mechanisms that establish neonatal enterocyte identity and coordinate their maturation remain poorly understood. Here we show that the transcription factors BLIMP1 and CASZ1 form a central switch in the timing of enterocyte maturation. While BLIMP1 marks and promotes neonatal enterocytes, CASZ1 is expressed in adult enterocytes and is required for the proper timing of their emergence. Genetic analysis indicates that these neonatal and adult factors are mutually repressive, forming a genetic switch underlying weaning. This regulatory mechanism also controls glucocorticoid-induced maturation, a process relevant to both normal weaning and the clinical use of corticosteroids to promote epithelial maturation in preterm infants. We show that corticosteroids act directly on the intestinal epithelium and require CASZ1 to induce maturation. Together, these findings reveal a transcriptional mechanism that coordinates intestinal epithelial identity with the changing nutritional demands of early life and mediates the response to maturation-promoting hormones.

## INTRODUCTION

The small intestine supports growth and survival by absorbing nutrients from diets that change dramatically between neonatal and adult life. While the absorptive intestinal epithelial cells (enterocytes) of newborns and adults share this critical core function, they utilize distinct mechanisms of nutrient uptake. While prior work has identified functional differences between neonatal and adult enterocytes, the mechanisms that control the transition from a neonatal enterocyte to an adult state remain unknown.

In the adult intestine, macromolecules are broken down into subunits within the lumen^1^, and specialized transporter proteins internalize these metabolites through the apical, microvilli-coated surface of enterocytes^2,3^. In contrast, neonatal enterocytes internalize dietary macromolecules whole and break them down in lysosomes^4–8^. This process, known as macromolecular uptake, is essential for neonatal growth^7^. These age-specific functions are also accompanied by distinct cellular morphologies^4,6,9,10^. Adult enterocytes have a well-developed brush border with a cytoskeleton-rich and vesicle-poor apical region, whereas neonatal enterocytes lack the dense apical cytoskeleton and instead have an apical region dominated by endocytic vesicles and large lysosomes^4,5,9,10^. Newborn rodents also undergo transcytosis of maternal milk-derived antibodies, providing immune support early in life^11^. Neonatal and adult enterocytes also differ in carbohydrate digestion: neonatal enterocytes highly express lactase to digest milk sugar, while adults utilize a suite of enzymes, including sucrase-isomaltase and trehalase, to break down diverse dietary saccharides^12–14^. Prior work has characterized these functional differences and identified the machinery that mediates endocytic uptake and lysosomal degradation in neonates^7,8^. However, it is not fully understood how these distinct enterocyte states are transcriptionally established.

The transcription factors BLIMP1 (gene: *Prdm1*), MAFB (gene: *MafB*), and cMAF (gene: *Maf*) are required for the neonatal state, and their loss impairs macromolecular uptake and growth^7,15,16^. Disrupting these neonatal transcription factors also induces the premature expression of adult enterocyte genes and acquisition of adult morphology^15,16^. This de-repression of the adult state suggests that antagonism between neonatal and adult programs may control postnatal intestinal maturation. However, the regulators that promote the adult state and their interplay with neonatal transcription factors are not known.

Beyond these intrinsic drivers of enterocyte identity, intestinal maturation is also regulated by systemic glucocorticoids^17–22^. In rodents, glucocorticoid levels increase during the third week of life ahead of weaning, coinciding with an increase in adult-specific enzymes^14^. Adrenalectomy delays the development of adult intestinal enzyme activity, and exogenous corticosteroids can induce precocious tissue maturation^17,19,23,24^. Together, these data support a role for these hormones in the neonatal-to-adult transition. Clinically, corticosteroids are administered to promote tissue maturation before premature birth, reducing rates of severe pulmonary disease, necrotizing enterocolitis, and death^25,26^. Although both “natural” maturation (i.e. occurring spontaneously ahead of weaning) and exogenous corticosteroid treatment induce functional changes within the intestinal epithelium, it remains unknown whether they operate through the same gene regulatory network.

In this study, we dissect the distinct transcriptional mechanisms that control neonatal enterocyte identity and the transition to adulthood. We identify the transcription factor CASZ1 as a key driver of the timing of the transition and reveal that CASZ1 and BLIMP1 act antagonistically. Ectopically extending BLIMP1 expression or ablating CASZ1 extends neonatal enterocyte identity and function. We also demonstrate that exogenous corticosteroids rely on CASZ1 to induce premature intestinal maturation. Together, this work provides mechanistic insight into the transcriptional programs that control postnatal intestinal maturation.

## RESULTS

### Molecular and Functional Maturation of Enterocytes Precedes Weaning

While mice are physically weaned from their dams at approximately 21 days of age, it was unclear if this process coincides with intestinal maturation on a cellular level. We therefore examined the expression dynamics of known neonatal transcription factors. Consistent with prior work^15,16,27^, we found that BLIMP1 is expressed broadly throughout the villi in early neonates (Fig. 1A-B). Between postnatal day 14 (P14) and P21, BLIMP1+ cells became progressively restricted to the tips of villi (Fig. 1A-B). During this same window of time, BLIMP1-negative cells first appeared at the villar base and then proceeded upward (Fig. 1A-B, Supp. Fig 1A). To determine how this progressive loss of BLIMP1+ cells impacted neonatal enterocyte function, we assessed macromolecular uptake by gavaging pups with fluorescently-labeled macromolecules^7,8^. The internalization of dietary macromolecules is a unique feature of neonatal enterocytes in the jejunum and ileum^7,8^, so this technique serves as a functional assay for neonatal enterocyte identification. Pups were given a bolus of a fluorescently-labeled macromolecule (Alexa Fluor 488-labeled dextran) via oral gavage and dissected three hours later. At P14, nearly all of the enterocytes showed evidence of macromolecular uptake, as demonstrated by internalized fluorescent macromolecules present throughout most of the villus (Fig. 1C, left). However, we consistently detected a region of cells (approximately 50 μm, typically 5-10 cells) at the base of P14 villi that did not internalize labeled macromolecules (Fig. 1C, left). Whether this reflects a region that does not have access to lumenal components, or whether this reflects a newly generated population of cells that has not acquired uptake activity remains unclear. When we performed this same analysis at P16, both the BLIMP1 domain and the uptake domain shifted farther up the villus (Fig. 1C right). Further, the rate of flux of the BLIMP1-negative and uptake-negative domains were very similar (Fig. 1D). This demonstrates a clear spatial and kinetic correlation between the loss of BLIMP1 and the loss of neonatal enterocyte function that precedes weaning.

**Figure 1:**
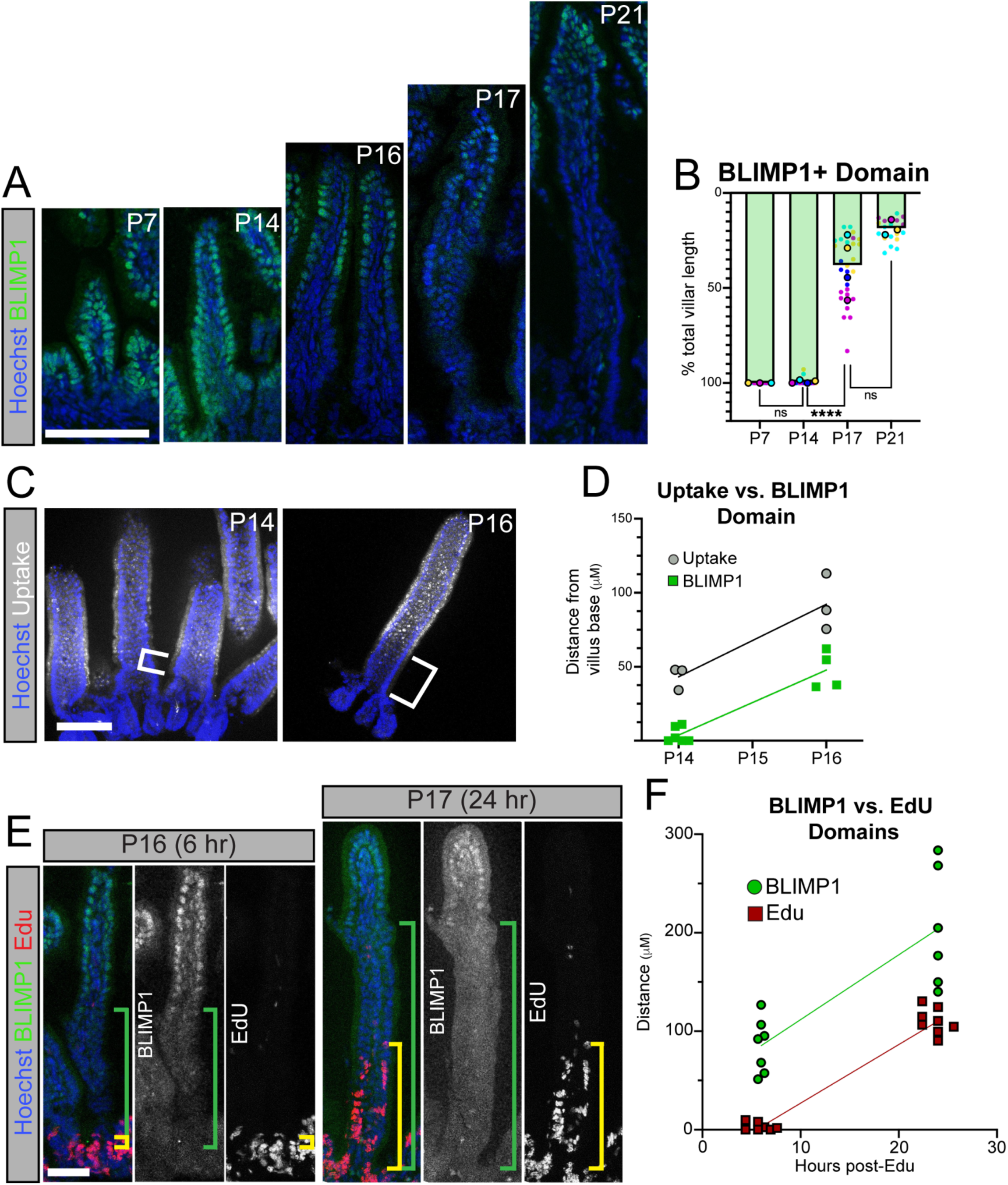
Molecular and functional maturation of enterocytes precedes weaning. **A-F**) Immunofluorescent (IF) detection (**A,C E**) and quantitation (**B**,**D, F**) of BLIMP1 (**A-B, E-F**), macromolecular uptake (**C-D**), and EdU incorporation (**F-G**) in postnatal mice. Superplot in **B** includes biological replicates (individual mice, circles with border) and the associated technical replicates (individual villi, color-matched open circles). Statistical significance in **B** determined with ordinary one-way ANOVA with Tukey’s multiple comparisons test. Distances in **D** and **F** measured from villus base. Brackets indicate the distance from the villus base to the macromolecular uptake domain (**C**, white), the BLIMP1+ domain (**E**, green), and the EdU+ domain (**E**, yellow). Nuclei marked in blue (Hoescht). Scale bars = 100 μm in **A** and **C**, 50 μm in **E**.

In the intestinal crypts, stem and transit-amplifying (TA) progenitor cells give rise to differentiated epithelial cells (including enterocytes), which then migrate progressively toward the tip of the villi before being shed into the epithelial lumen^28^. This flux of differentiated epithelial cells from the base of the villus to the tip occurs over 3-5 days in mature animals^29^. Given the progressive loss of BLIMP1 and neonatal enterocyte function ahead of weaning, we investigated if the loss of BLIMP1+ cells was due to epithelial flux. In this model, a switch in enterocyte progenitors begins producing adult cells to replace the BLIMP1+ neonatal cells. Alternatively, the observed loss in BLIMP1 could reflect active repression in cells already expressing BLIMP1. To differentiate between these two possibilities, we performed EdU pulse-chase assays. We injected mice with EdU at P16 and collected mice 6 and 24 hours later. We then compared the position of EdU-labeled cells with the position of BLIMP1+ cells. At 6 hours post-EdU injection, we confirmed that the pulse-labeled BLIMP1-negative cells were positioned at the base of the villus (Fig. 1E, left). When we assessed the BLIMP1 and EdU domains 24 hours post-EdU injection, we found that the rates of movement up the villi paralleled each other (Fig. 1E, right; Fig, 1F). This suggests that the enterocyte progenitor cells in the crypt switch from producing neonatal enterocytes to mature enterocytes, and this wave of mature enterocytes replaces neonatal cells through epithelial flux. Thus, both the molecular and physiological characteristics of neonatal enterocytes are lost in a switch-like manner, and this precedes the physical weaning of the pups.

In order to gain a more mechanistic understanding of the neonatal-to-adult transition, we first wanted to define the neonatal transcriptional program driven by BLIMP1. Prior work identified 254 BLIMP1 regulated genes using microarray analysis^16^, and we sought to expand on this list using bulk RNA sequencing (RNA-seq) on *Prdm1* WT and conditional knockout (*Prdm1* cKO: *Villin^Cre/+^;Prdm1^fl/fl^*) intestinal epithelial cells (Supp. Fig. 1B). Our RNA-seq analysis identified over 5,500 differentially expressed genes, including nearly 3,000 upregulated genes and approximately 2,600 downregulated genes (Supp. File 1, Supp. Fig 1C). Consistent with the existing *Prdm1* cKO microarray data^16^, we detected a significant upregulation in adult-specific enzymes such as sucrase-isomaltase (*Sis*) (Supp. Fig. 1D), trehalase (*Treh*), adenosine deaminase (*Ada*), and arginase 2 (*Arg2*) (Supp. File 1). However, we also detected significant upregulation of many other genes required for adult enterocyte function, including apical amino acid transport (*Slc1a1, Slc15a1, Slc7a15*) (Supp. Fig. 1E-G), a process utilized by enterocytes to internalize digested lumenal peptides^30–32^. We also detected a significant upregulation in the adult-specific enzyme maltase-glucoamylase (*Mgam*) required for carbohydrate metabolism^33^. The premature expression of these adult-specific genes supports the existing model that BLIMP1 suppresses the adult state in the neonatal epithelium^15,16^. Down-regulated genes included those encoding endolysosomal machinery (*Ctsl*, *Lgmn*, and *Dab2*) (Supp. Fig. 1H), which are also regulated by MAFB/cMAF and are associated with neonate-specific macromolecular uptake^7,8^. Additionally, there was significant downregulation of the immunoglobulin transporter (*Fcgrt*) in *Prdm1* cKO neonates (Supp. File 1). To determine if these changes in gene expression impacted neonatal epithelial cell metabolism, we performed metabolomic analysis of WT and *Prdm1* cKO epithelial cells. We also analyzed *MafB;cMaf* cKO intestinal epithelial cells, as both *Prdm1* cKO and *MafB;cMaf* cKO mice display neonatal growth defects^7^. Conditional deletion of *Prdm1* or *MafB/cMaf* reduced the concentration of all amino acids in epithelial cells (Supp. Fig. 1I, Supp. File 2). This dramatic decrease in amino acid levels likely explains, at least in part, the severe growth defects observed in *Prdm1* cKO and *MafB/cMaf* cKO neonates^7,15,16^. Further, it demonstrates that upregulation of the adult enterocyte program is not sufficient for protein/amino acid uptake in neonates. Together, these data indicate that BLIMP1 is required to activate multiple key functional aspects of neonate-specific cell identity, including both neonatal nutrient absorption and maternal-derived immune support.

### CASZ1 expression coincides with the transition to an adult state

While transcription factors have been identified that direct neonatal enterocyte identity, it remains unknown whether the adult state is induced by adult-specific factors, or whether the adult state is the default in the absence of neonatal factors. To investigate this question, we first generated an adult-specific epithelial gene list. To accomplish this, we compared our *Prdm1* cKO RNA-seq dataset to other gene expression datasets that all share an adult-like phenotype, in order to identify which genes constitute the “core” of the adult functional state. These additional phenotypically adult datasets included RNA-seq of natural maturation^34^ (P6 vs P36), RNA-seq of an additional pre-mature maturation model^7^ (*MafB;cMaf* cKO: *Villin^Cre/+^;MafB^fl/fl^;cMaf^fl/fl^*), and scRNA-seq analysis of TA progenitor cells before (P10) and after (P14) the switch from producing neonatal to adult enterocytes (Y.W. and K.S. – Manuscript in preparation). This analysis identified an adult core (Fig. 2A, Supp. File 3) of 853 genes that were detected in at least three out of the four datasets. Pathway analysis identified a strong metabolic signature associated with the adult phenotype, featuring genes required for sugar metabolism and lipid breakdown (Fig 2B). In parallel, we generated a neonatal core list that included 1,078 neonate-specific genes (Fig. 2C, Supp. File 3). The neonatal core included a strong endolysosomal signature (Fig. 2D), consistent with the macromolecular uptake mechanism employed by these cells. This analysis identifies genes that differentiate neonatal from adult epithelial cells and elucidates a coordinated metabolic program that allows epithelial cells to match their age-specific diet.

**Figure 2:**
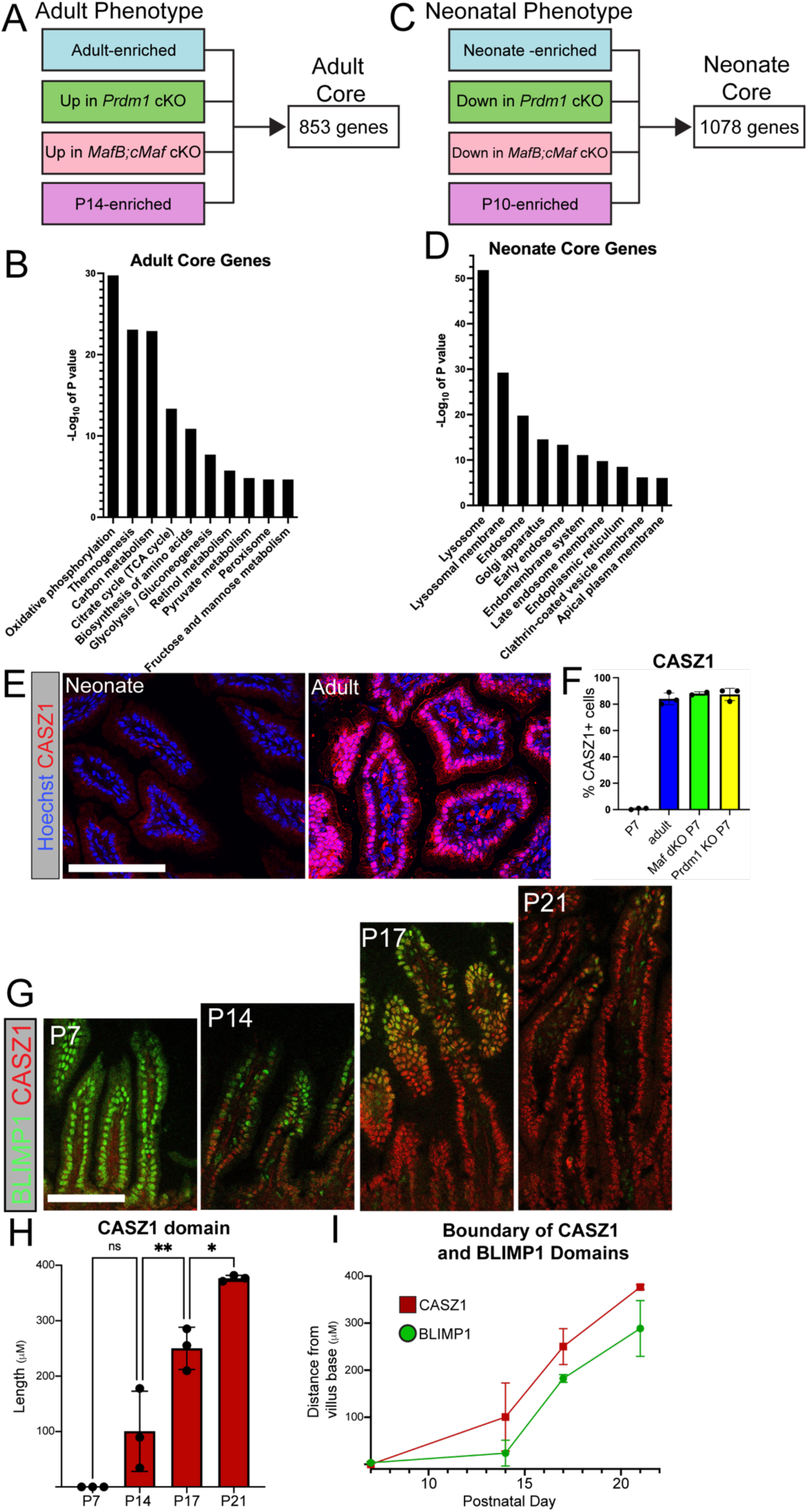
Identification of adult enterocyte-specific markers. **A-D**) Multi-dataset analysis identifying “adult core” (**A-B**) and “neonate core” (**C-D**) genes in intestinal epithelial cells. Source datasets include neonate vs. adult analysis^1^, WT vs. *Prdm1* cKO neonate analysis (this paper), WT vs. *MafB;cMaf* cKO neonate analysis^2^, and P10 vs. P14 TA cell analysis (Y.W. and K.S. – Manuscript in preparation). Pathway analysis in **B** and **D** determined by KEGG and GO analysis, respectively. Terms displayed represent a selected set of significantly enriched pathways. **E-I**) IF antibody detection (**E,G**) and quantitation (**F, H, I**) of CASZ1 at the indicated ages and in the indicated mouse models. Quantitation in **F** depicts the percentage of cells expressing CASZ1 within 100 μm of the villus base. Length measurements in **H** and **I** measured from the villus base. Additional antibodies include BLIMP1 (green, **G**). Statistical significance in **H** determined with ordinary one-way ANOVA with Tukey’s multiple comparisons test. Each point in **H** represents the average value for an individual animal. Each point in I represents the mean value across a minimum of three animals. All scale bars = 100 μm.

We sought to refine these results to identify transcriptional markers of the adult phenotype. By interrogating our adult core gene list and isolating known transcriptional regulators, we extracted two candidates of interest: ID3 and CASZ1. While these transcription factors have established roles in regulating the development of other cell types^35–37^, their role in the small intestinal epithelium is unknown. Both *Id3* and *Casz1* are expressed at higher levels in adults compared to neonates^34^ (Supp. Fig. 2A), and become upregulated in *Prdm1* cKO neonates (Supp Fig. 2B). We validated our gene expression analysis by performing immunofluorescence analysis (IF) which demonstrated that CASZ1 and ID3 expression increased in adults, *Prdm1* cKO neonates, and *MafB;cMaf* cKO neonates compared to WT neonate controls (Fig. 2E-F, Supp. Fig. 2C). However, in adult mice epithelial CASZ1 is expressed along the entire length if the villi (Fig. 2E), whereas epithelial ID3 remains restricted to the upper regions of the crypt and the very base of the villi (Supp. Fig. 2C). Together, these data indicate that both CASZ1 and ID3 are specific to the adult intestinal epithelium.

Enterocyte maturation occurs between P14-P21, so we assessed the expression dynamics of ID3 and CASZ1 across this period. We confirmed by IF that ID3 is initially absent in the P7 neonatal epithelium, although ID3+ mesenchymal cells were detected at all stages (Supp. Fig. 2D-E). By P14 we detected clear epithelial expression of ID3 in the upper region of the crypt (Supp. Fig 2D-E). While the abundance of ID3+ cells increased from P14 to adulthood (Supp Fig. 2E), it remained restricted to the top of crypts and the villar base, and was excluded from the upper villus and the bottom of the crypt. This pattern is consistent with TA-specific expression, and loss of gene expression after villar commitment. In addition to an increase in ID3+ cell number, we also detected an increase in ID3 nuclear fluorescence intensity levels between P17 and weaning age (Supp Fig. 2F).

CASZ1 was also absent from the early neonatal epithelium at P7 (Fig. 2G-H) with expression at the base of the villus becoming apparent at P14. Its expression throughout the villus increased over time, moving up in a wave. The onset and dynamics of CASZ1 expression opposed loss of BLIMP1 expression (Fig. 1A). We therefore compared the expression of these transcription factors directly and found an opposing expression pattern (Fig 2G,I). Together, these data demonstrate a molecular switch in transcription factor expression during intestinal maturation, that coincides with the neonatal-to-adult transition.

### BLIMP1 is sufficient to maintain neonatal intestinal identity

If this transcriptional switch from a neonatal-to-adult cell state is required for maturation, we predicted that forced maintenance of *Prdm1* expression would extend the neonatal state. To test this, we used a mouse line in which ectopic *Prdm1* expression can be activated following CRE-mediated recombination of an upstream loxP-STOP-loxP sequence (*ROSA26:LSL-Prdm1*)^38^. We crossed this mouse line to the *Villin^CreER^*line (*Prdm1^Tg^: Villin^CreER/+^; ROSA26:LSL-Prdm1*) to inducibly drive ectopic *Prdm1* expression in the intestinal epithelium (Fig. 3A), and confirmed strong BLIMP1 expression throughout the intestinal epithelium upon tamoxifen administration (Fig. 3B). We induced ectopic *Prdm1* expression at P12 (before the neonatal-to-adult transition), collected mice at P16, and stained for CASZ1 as a marker of adult enterocytes. At this stage, CASZ1 marked cells at the base of the villus in control mice, but this expression was almost completely blocked in ectopic *Prdm1*-expressing cells (Fig. 3C-D). When we assessed ID3 expression in *Prdm1^Tg^* mice, we found that a majority of *Prdm1^Tg^* mice expressed ID3 in the upper crypt, and did not show any significant change in abundance between WT and *Prdm1^Tg^* mice (Supp. Fig. 3A-B). Therefore, BLIMP1’s antagonism of the adult transcription factor candidates is specific to CASZ1.

**Figure 3:**
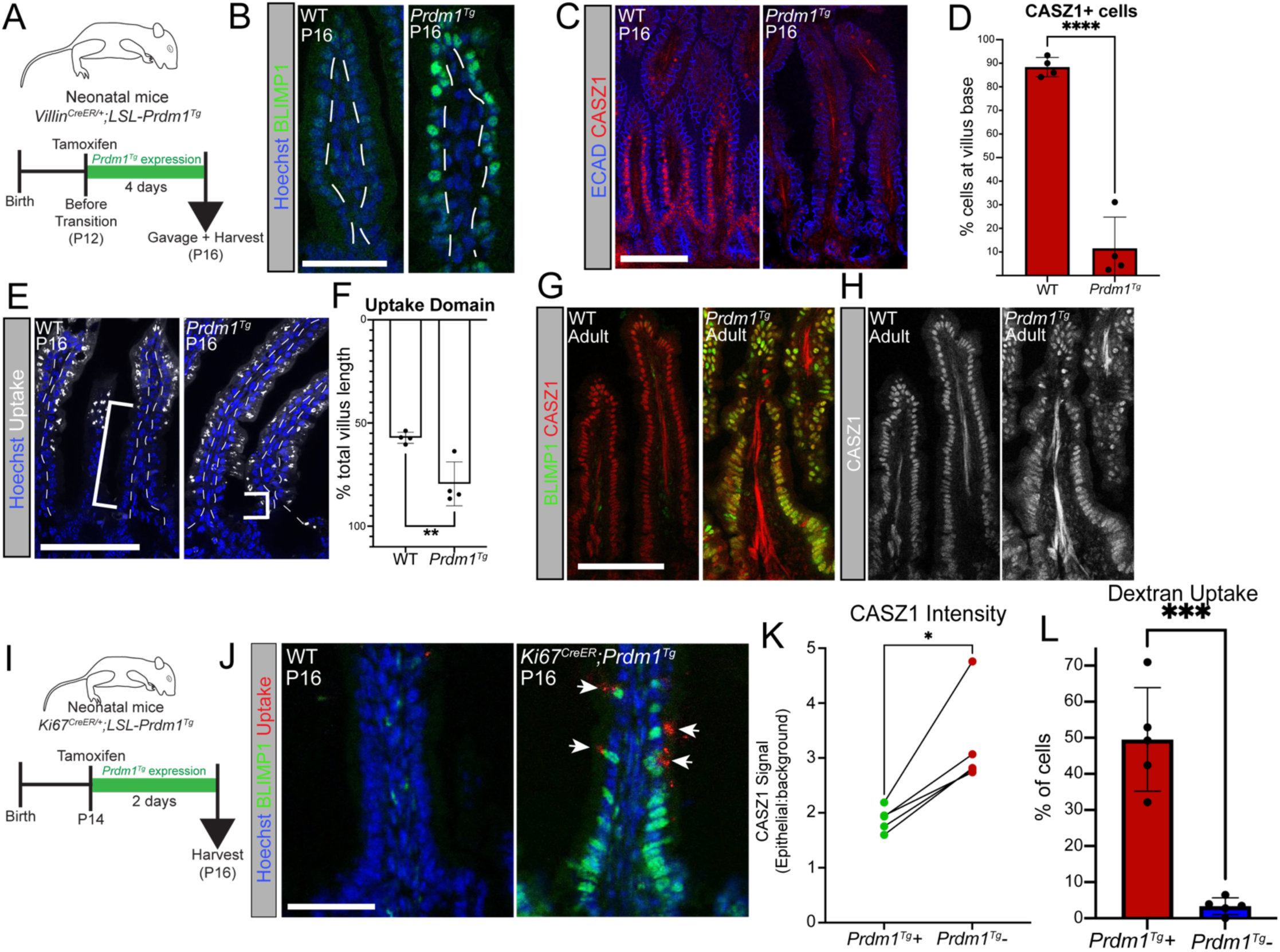
BLIMP1 is sufficient to maintain neonatal intestinal identity. **A-F)** Ectopic expression of *Prdm1* via the *Villin^CreER/+;^ROSA26:LSL-Prdm1* (*Prdm1^Tg^*) from P12 to P16. Cartoon in **A** depicts experimental strategy. IF analysis includes antibody detection of BLIMP1 (green, **B**), CASZ1 (red, **C**), ECAD (blue, **C**), as well as detection of macromolecular uptake (Alexa Fluor 488-labeled dextran, white, **E**) by neonatal enterocytes. Quantitation measures the percentage of cells that express CASZ1 (**D**) and the percentage of the total villus that has undergone macromolecular uptake (**F**). **G-H**) Analysis of BLIMP1 (green) and CASZ1 (red in **G**, grayscale in **H**) expression following ectopic *Prdm1* expression in mature animals. **I-L**) Analysis of ectopic *Prdm1* expression (green) and macromolecular uptake (Alexa Fluor 488-labeled dextran, red) in *Ki67^CreER/+^;ROSA26:LSL-Prdm1* mice (*Ki67^CreER^;Prdm1^Tg^*), as determined by IF analysis. Experimental strategy depicted in **I**. Quantitation in **K-L** depicts the percentage of BLIMP1+ cells expressing CASZ1 (**K**) or undergoing macromolecular uptake (**L**) in *Ki67^CreER^;Prdm1^Tg^* mice, compared to BLIMP1-neighboring cells. Statistical significance in K determined by paired t-test. For all other quantitation, statistical significance determined by unpaired t-test. For all quantitation, each point represents an independent animal. Nuclei in **B**, **E**, and **J** marked in blue (Hoescht). Scale bars = 50 μm in **B** and **J**, 100 μm in **C**,**E**, and **G**.

To evaluate whether prolonged expression of BLIMP1 affected neonatal enterocyte function, we performed gavage assays. Consistent with our previous experiments, WT P16 pups are in the midst of the neonatal-to-adult transition, and therefore only maintain macromolecular uptake in the upper region of the villi (Fig. 3E, left). However, P16 *Prdm1^Tg^* mice maintained macromolecular uptake throughout the length of the villus (except at the villar base, in which we never detect uptake), consistent with the loss of adult markers (Fig. 3E right, 3F). This indicates that maintaining BLIMP1 expression throughout the neonatal-to-adult transition keeps enterocytes locked in a neonatal functional state. Unfortunately, *Prdm1^Tg^* mice typically do not survive more than a week after ectopic *Prdm1* activation, precluding longer-term analysis of enterocyte identity in mice that maintain *Prdm1^Tg^* expression further into adulthood. While we suspect that this lethality is due to non-intestinal phenotypes, the ultimate cause of this lethality remains unknown.

To determine if BLIMP1 expression is sufficient to convert adult enterocytes back into a neonatal state, we induced ectopic *Prdm1* expression in fully mature adult mice (Supp Fig. 3C). While ectopic *Prdm1* expression starting at P12 nearly eliminated CASZ1 in P16 mice (Fig. 3C-D), the majority of adult epithelial cells maintain CASZ1 when BLIMP1 is expressed (Fig. 3G-H). The adult *Prdm1^Tg^*mice also featured a substantial number of cells that co-express both BLIMP1 and CASZ1 (Supp. Fig. 3D-E), which was rarely observed during natural maturation (Fig. 2G). Although there is a slight reduction in CASZ1+ cells in *Prdm1^Tg^*adults (Fig. 3H), the persistence of this adult transcription factor suggests that BLIMP1 largely loses the ability to suppress the adult program once maturation is complete. Further, *Prdm1^Tg^*adults also fail to activate the expression of neonatal genes (e.g. *Lgmn*) (Supp Fig. 3F). Together, these data indicate that once epithelial cells have completed the neonatal-to-adult transition and reach maturity, they become refractory to BLIMP1’s neonate-inducing activity.

To determine if the neonatal-promoting effect of BLIMP1 is cell-autonomous, we took advantage of the *Ki67^CreER^* mouse line^39^ (Fig. 3I). The *Ki67^CreER^* allele induces recombination in a subset of TA cells, leading to a mosaic pattern of CRE-recombined cells in the villi^40^. In *Ki67^CreER^* mice, maintenance of macromolecular uptake is limited to cells that express ectopic *Prdm1*, while *Prdm1-*negative neighboring cells express higher levels of CASZ1 and do not perform macromolecular uptake (Fig. 3J-L). Thus, this model demonstrates that ectopic BLIMP1 maintains neonatal function in a cell-autonomous manner. Further, these cell-autonomous phenotypes rule out the possibility that our *Villin^CreER^; Prdm1^Tg^* phenotypes (Fig. 3C-F) are merely due to a tissue-wide postnatal developmental delay.

### CASZ1 is required for proper timing of intestinal maturation

Our analysis of the neonatal-to-adult transition identified two transcription factors associated with the adult enterocyte cell state: CASZ1 and ID3. CASZ1 demonstrated an opposing expression pattern with BLIMP1 (Fig. 2G), and was suppressed by maintained *Prdm1* expression (Fig. 3C). Given the BLIMP1-driven antagonism of CASZ1, we next investigated whether CASZ1 affected BLIMP1 expression. To test this, we conditionally deleted *Casz1* (*Casz1^fl/fl^*)^41^ from the intestinal epithelium (*Casz1* cKO: *Villin^Cre/+^;Casz1^fl/fl^*) and verified loss of expression via immunofluorescence (Fig 4A-B). We examined pups at P16, a time at which WT littermate controls have begun the transition to the adult phenotype and BLIMP1 is lost at the base of the villi. However, *Casz1* cKO pups maintain BLIMP1 expression at the base of villi (Fig. 4A, C). In addition, the macromolecular uptake domain extended further toward the villar base in *Casz1* cKO pups (Fig. 4D-E), approaching the maximum domain of macromolecular uptake observed in P14 pups (down to 50 μm, Fig. 1C-D). This indicates that the loss of *Casz1* prolongs neonatal enterocyte function. To determine if the neonatal-to-adult transition is delayed or eliminated in *Casz1* cKO mice, we collected *Casz1* cKO mice at P20 and P25. At P20, *Casz1* cKO mice begin to lose BLIMP1+ cells at the villar base, and by P25 the domain of BLIMP1+ is indistinguishable from WT littermate controls (Fig. 4F-G). Thus, our data indicate that CASZ1 represses both BLIMP1 expression and neonatal enterocyte function during the neonatal-to-adult transition. However, while CASZ1 is required for the proper timing of gut maturation, a delayed neonatal-to-adult transition can still occur independently of CASZ1. Adult *Casz1* cKO animals displayed no defects in total mature body weight (Supp Fig. 4A), suggesting that the absence of *Casz1* does not lead to lasting defects in gross body development. Together, our data demonstrate that the intestinal epithelium has compensatory mechanisms to ensure tissue maturation in the absence of *Casz1*.

**Figure 4:**
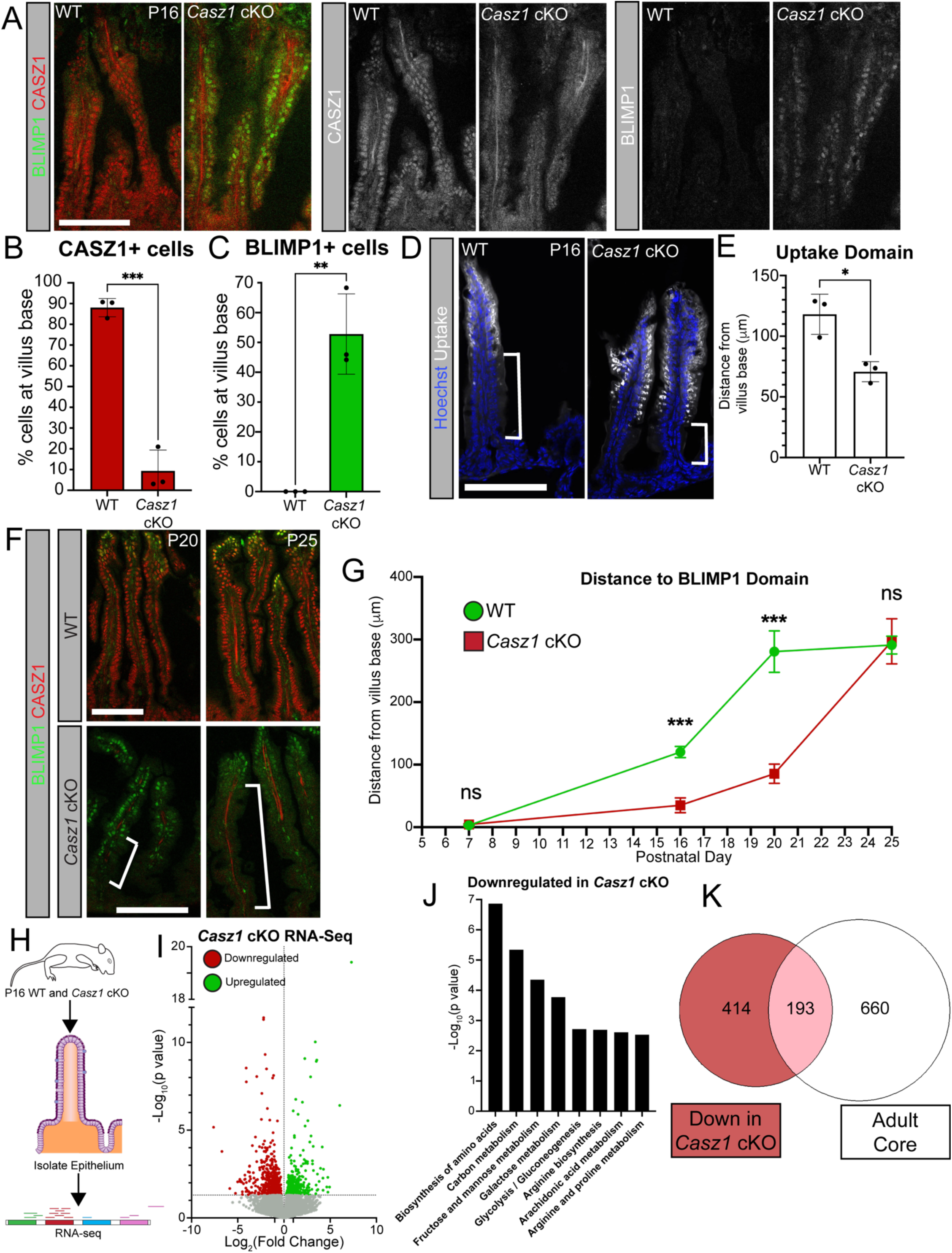
CASZ1 is required for proper timing of intestinal maturation. **A-G)** IF analysis of postnatal *Casz1* cKO mice. Analysis includes antibody detection of CASZ1 (**A**, **F**, red), BLIMP1 (**A**, **F**, green), and detection of macromolecular uptake (Alexa Fluor 488-labeled dextran, white, **D**). Grayscale images in **A** display single channel images of CASZ1 and BLIMP1 staining, respectively. Quantitation in **B-C** represents the percentage of cells within 75 μm of the villus base that express the indicated transcription factor. Quantitation in **E** and **G** measures the distance from the villus base to the indicated signal (depicted as white brackets in **D** and **F**). For all quantitation **B-G**, statistical significance is determined unpaired t-test. All experiments used a minimum of three animals at each timepoint (each point in **B**, **C**, **E** represents an individual animal’s average value, points in **G** represent the mean value across all biological replicates). Scale bars for IF images = 100 μm. Nuclei in **D** marked in blue (Hoescht). **H-K)** Gene expression analysis of P16 WT vs. *Casz1* cKO intestinal epithelial cells, as determined by RNA-seq. Cartoon in **H** depicts experimental strategy. Volcano plot in **I** depicts genes that are significantly upregulated (green), downregulated (red), or not significantly changed (gray). Dotted lines indicate p = 0.05 (Y axis) and a Fold change = 1 (X axis). Pathway analysis displayed in **J** includes a selected list of significantly downregulated gene pathways in *Casz1* cKO mice, as determined by KEGG analysis. Venn diagram in **K** depicts the unique and shared genes that are downregulated in *Casz1* cKO mice (red) and present in the adult core gene list.

In contrast to loss of *Casz1*, loss of *Id3* did not cause defects in intestinal maturation. Germline *Id3^-/-^* mice ^36^ are viable, born at mendelian ratios, and show clear loss of ID3 staining in the crypts (Supp. Fig.4B). The absence of *Id3* had no impact on macromolecular uptake (Supp Fig. 4C-D), nor BLIMP1/CASZ1 expression (Supp. Fig. 4E-F), indicating that the progressive loss of the neonatal program and adoption of the adult program occurs normally in the absence of ID3. Therefore, while ID3 and CASZ1 are both upregulated during the neonatal-to-adult transition, only CASZ1 appears to be functionally required for the proper timing of intestinal maturation.

As CASZ1 is required for timely epithelial maturation, we next wanted to identify CASZ1 regulated genes to determine the extent of its effects on the transcriptome. We isolated epithelial cells from P16 WT and *Casz1* cKO mice (Fig. 4H), and analyzed differential gene expression by RNA-seq (Supp. File 4). We determined that 960 genes were differentially expressed in *Casz1* cKO P16 intestinal epithelial cells compared to WT littermate controls, with roughly twice as many downregulated genes (607) as upregulated genes (354) (Fig. 4I). The relatively modest number of differentially expressed genes is likely due to a mixed population of both neonatal and adult-like epithelial cells at P16. Nevertheless, we detected patterns of differential gene expression that provide insight into the role of CASZ1 during the neonatal-to-adult transition. Specifically, we found a significant decrease of adult-specific gene expression in *Casz1* cKO mice. Pathway analysis of the downregulated genes in *Casz1* cKO mice revealed changes in genes regulating metabolic processes. Specifically, we found downregulation of amino acid and sugar metabolism pathways that are unique to adult enterocytes (Fig. 4J). By directly comparing the *Casz1* cKO downregulated gene list to the adult core genes (Fig. 4K), we determined that CASZ1 activates, directly or indirectly, 193 of the adult core genes. These CASZ1 target genes include apical transporter proteins (*Slc2a5, Slc22a4*), brush border enzymes (*Treh, Sis, Mgam*), and genes required for amino acid (*Got1*, *Arg2*) and sugar metabolism (*Gpi1*, *AldoA*) (Supp File 4). Thus, these data indicate that CASZ1 drives the neonatal-to-adult transition by activating an adult-specific metabolic program.

### Glucocorticoid-driven intestinal maturation requires CASZ1 activation and BLIMP1 repression

While our data identifies intrinsic factors that direct epithelial maturation, extracellular cues can also impact epithelial cell function. The adult epithelial cell state can be induced prematurely by treatment with glucocorticoid receptor agonists, such as dexamethasone (dex)^17,42^. Early postnatal administration of glucocorticoids in rodents induces changes in gene expression and enzyme activity consistent with premature maturation^17,21,43^. Additionally, removal of the adrenal gland, a major source of systemic glucocorticoids, delays enterocyte maturation^19,23,24^. However, the transcriptional mechanisms underlying hormone-induced maturation, and the degree of overlap with the natural maturation mechanisms, are unknown.

To investigate this, we treated neonatal mice daily with dex from P4 to P6 to induce premature maturation, and collected them on P7 to assess neonatal vs. adult gene expression (Fig. 5A). Dex treatment resulted in the loss of BLIMP1 and upregulation of CASZ1 expression (Fig. 5B-D), demonstrating that the core transcriptional drivers of natural maturation are altered by glucocorticoid signaling. To globally define the dex-driven transcriptional program, we performed RNA-seq on isolated intestinal epithelial cells from vehicle and dex-treated P7 mice. Dex treatment led to broad transcriptional changes in the neonatal epithelium, with over 5,500 differentially expressed genes (Supp File 5). To determine how dex-induced maturation compared to other models of maturation, we compared the dex-induced differentially expressed genes to our adult and neonatal core gene lists. Neonates treated with dex upregulate ∼60% of the adult core genes and downregulate ∼61% of the neonatal core genes (Supp. Fig 5A-C). To compare dex-induced changes to gene expression to intrinsic models of premature maturation, we also compared the dex RNA-seq dataset to the *Prdm1* cKO RNA-seq dataset. Similar to what we observed when comparing dex-treated neonates to the adult core (Supp. Fig 5A), only a subset of *Prdm1* cKO differentially expressed genes were significantly changed following dex (Supp. Fig 5D). However, the genes that are shared between both datasets show a strong correlation in gene expression changes (Supp. Fig. 5E). These data demonstrate that dex-induced maturation shows many similarities with natural maturation, including the core transcriptional regulator switch.

**Figure 5:**
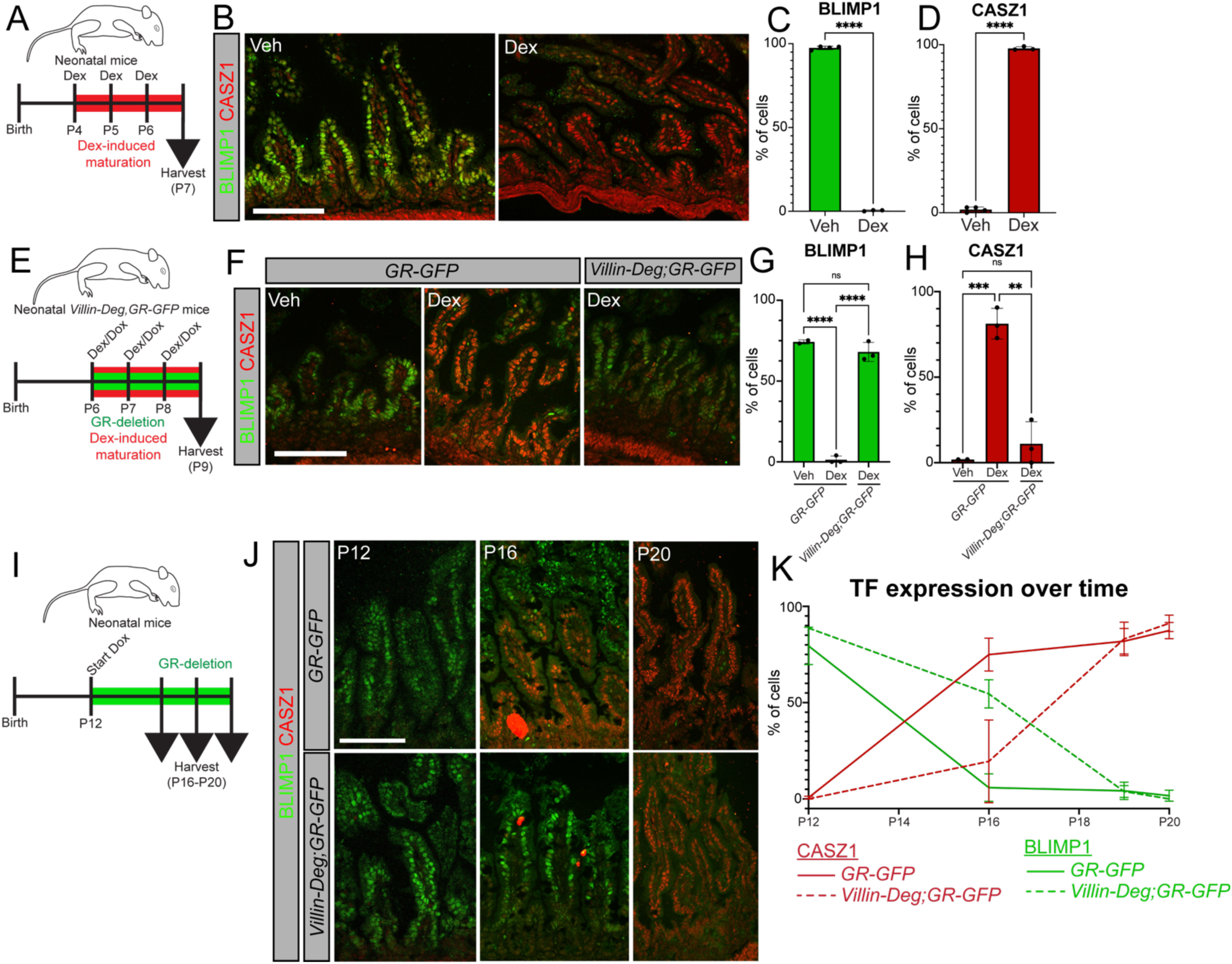
Glucocorticoid stimulation of the intestinal epithelium drives CASZ1 expression and BLIMP1 repression. **A)** Cartoon depicting experimental strategy: characterization of BLIMP1/CASZ1 expression following three days (P4, P5, P6) of dexamethasone (dex) i.p. injections (dosage = 0.4 mg/kg, diluted in H2O). Vehicle-treated animals received an equivalent volume of H2O only. **B-D)** IF detection (B) and quantitation (C-D) of BLIMP1 (green) and CASZ1 (red) following three days of dex injections. **E)** Cartoon depicting experimental strategy: characterization of BLIMP1/CASZ1 expression in P9 *Villin-Deg;GR-GFP* mice following three days of dex injections. Control animals are littermate controls that lack a functional degron system (lacking Villin-rtTA or Tre-Degron), but still received doxycycline. **F-H)** IF detection (F) and quantitation (G-H) of BLIMP1/CASZ1 in the indicated mice. **I)** Cartoon depicting experimental strategy: determining the timing of natural maturation in mice lacking GR. **J)** IF detection of BLIMP1 (green) and CASZ1 (red) over time in *Villin-Deg;GR-GFP* mice (and degron-negative littermate controls). **K)** Summary plot of CASZ1 (red) and BLIMP1 (green) expression dynamics over time in *Villin-Deg;GR-GFP* mice (dashed lines) and degron-negative littermate controls (solid line). All quantitation plots represent the percentage of cells within 75 μm of the villus base that express the indicated transcription factor. At each timepoint, statistical significance was determined by unpaired t-test. Scale bars = 100 μm.

We next wanted to determine whether dex treatment led to permanent changes in the postnatal intestine. We therefore treated neonatal mice with three consecutive days of dex at P1, P2, and P3, and collected them at P11 (Supp. Fig. 5F). We also collected a cohort of mice at P4, and confirmed that early neonates were still responsive to dex, leading to suppression of BLIMP1 and activation of CASZ1 (Supp Fig. 5G, left). However, after an 8 day “washout” following dex treatment, P11 mice were largely comparable to vehicle-treated mice, displaying only a slight shift in the BLIMP1+ domain and a complete absence of CASZ1 (Supp. Fig. 5G-I). This indicates that the dramatic dex-dependent changes are both temporary and reversible.

While these data indicate that dex drives epithelial maturation, the drug is administered systemically in our experiments. Similarly, prior investigation into the intestinal role of these hormones used techniques to stimulate glucocorticoid activity in all cell types^43^, or impaired systemic corticosteroid synthesis through adrenalectomy^19,23,24^. Therefore, it was not known whether dex-induced intestinal maturation is a direct or secondary effect on the epithelium. To address this, we took advantage of a recently developed *in vivo* degron system, to selectively degrade GFP-tagged proteins of interest in a conditional, doxycycline-inducible manner^44^. Specifically, we crossed a *Villin^rtTA^* allele^45^ to a *TRE-Degron^GFP^* allele in order to selectively target GFP-tagged proteins for degradation in the intestinal epithelium^44^. We then incorporated a homozygous GFP-tagged Glucocorticoid Receptor allele (*GR-GFP*^46^), generating a *Villin^rtTA^;TRE-Degron^GFP^;GR-GFP/GR-GFP* mouse line (*Villin-Deg,GR-*GFP) that can conditionally and inducibly deplete GR in intestinal epithelial cells. We first validated that this mouse model effectively depletes GR-GFP from intestinal epithelial cells *Villin-Deg,GR-GFP* mice, and that epithelial GR-GFP is maintained in control littermates (Supp. Fig. 5J). Importantly, GR-GFP expression is maintained in underlying mesenchymal cells (Supp. Fig. 5J), indicating that this mouse model selectively depletes GR-GFP from the intestinal epithelium. To determine if dex-induced epithelial maturation is due directly to glucocorticoid signaling in the epithelium, we treated *Villin-Deg,GR-GFP* mice with both dex (to active GR) and doxycycline (to degrade GR) daily starting at P6 (Fig. 5E). We collected mice at P9, and evaluated neonatal and adult gene expression. In control littermates dex caused a loss of BLIMP1 and premature expression of CASZ1 (Fig. 5F-H). In contrast, *Villin-Deg,GR-GFP* mice failed to respond to dex and maintained BLIMP1 expression, with no upregulation of CASZ1 (Fig. 5F-H). Thus, these data indicate that the intestinal epithelium undergoes dex-induced maturation due to direct glucocorticoid signaling via epithelial GR.

Beyond exogenous hormone-induced maturation, prior work has indicated that impairing systemic corticosteroid levels via adrenalectomy disrupts natural intestinal maturation^19,23,24^. To determine if this phenotype is due to direct epithelial glucocorticoid signaling, we administered doxycycline to *Villin-Deg,GR-GFP* mice ahead of the neonatal-to-adult transition (P12), and maintained mice on doxycycline until the collection of tissues at multiple postnatal timepoints (Fig. 5I). As a control, we collected a cohort of mice at P12, and confirmed that depleting GR-GFP ahead of the neonatal transition had no effect on transcription factor expression, as the presence of BLIMP1 and absence of CASZ1 was consistent in both *Villin-Deg,GR-GFP* and GR-GFP control pups (Fig. 5J-K). At P16, *Villin-Deg,GR-GFP* mice maintained BLIMP1 expression at the villar base and lacked CASZ1, indicating that epithelial maturation was delayed (Fig. 5J-K). However, by P20 *Villin-Deg,GR-GFP* and control littermates had comparable expression of CASZ1 throughout the villi, and BLIMP1 was largely absent (Fig. 5J-K). These data reveal that endogenous glucocorticoid signaling is required for proper timing of the neonatal-to-adult transition, but that GR-independent mechanisms are eventually able to drive tissue maturation.

Our data demonstrate that hormone-induced maturation activates endogenous adult transcription factors (CASZ1) and suppresses neonatal transcription factors (BLIMP1). However, it doesn’t address whether those factors are required for glucocorticoid-induced maturation. We therefore manipulated this process through two different models: forced expression of *Prdm1*, and conditional deletion of *Casz1*. To achieve the former, we activated *Prdm1^Tg^* expression in P7 mice, followed by three consecutive days of dex treatment (Fig. 6A). As previously demonstrated, WT pups downregulated BLIMP1 and expressed CASZ1 in response to dex-treatment (Fig. 6B, left). However, *Prdm1^Tg^* mice maintained strong ectopic BLIMP1 expression and maintained neonatal enterocyte function (macromolecular uptake) in the presence of dex, and had no expression of CASZ1 (Fig. 6B, right, Fig. 6C-E). To test this same question in a distinct, cell-autonomous model, we performed a parallel experiment in *Ki67^CreER^;Prdm1^Tg^* mice (Fig. 6F). *Ki67^CreER^;Prdm1^Tg^-*expressing cells treated with dex suppress CASZ1 and maintain macromolecular uptake, whereas *Prdm1^Tg^*-negative neighbors respond to dex by expressing CASZ1 and shutting down macromolecular uptake (Fig. 6G-I). To determine if CASZ1 is required for dex-induced maturation, we treated *Casz1* cKO mice with dex from P8-P10 and collected intestines at P11 (Fig. 6J). In the absence of *Casz1*, dex was unable to suppress BLIMP1 expression, leading to BLIMP1 expression throughout the villi, a failure to express CASZ1, and the maintenance of macromolecular uptake (Fig. 6K-O). Together, these data support a model in which glucocorticoids act through altering the relative expression of BLIMP1 and CASZ1, who mutually antagonize one another.

**Figure 6:**
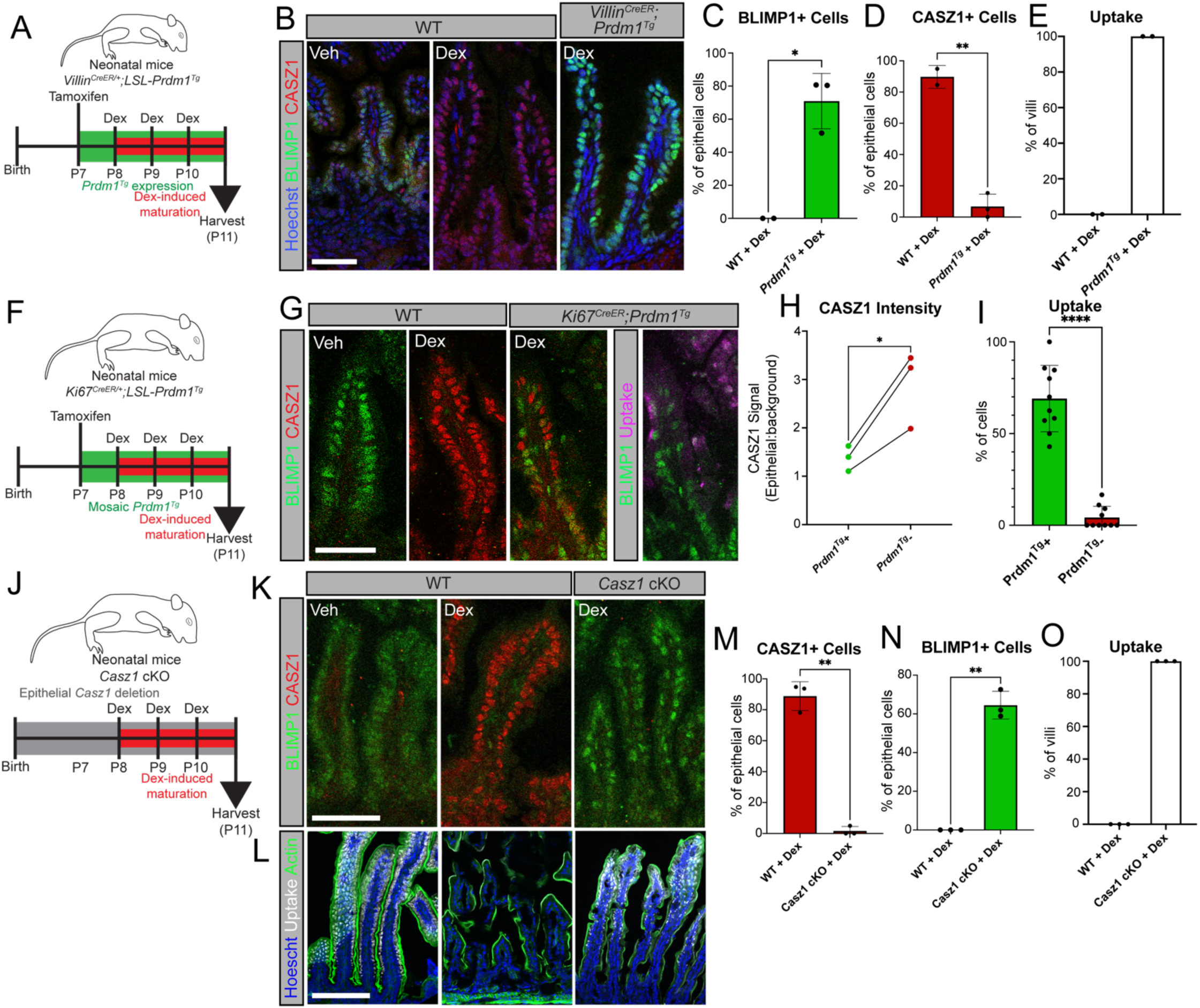
CASZ1 is required for glucocorticoid-induced maturation. **A)** Cartoon depicting experimental strategy: characterization of CASZ1/BLIMP1 expression in *Villin^CreER^;Prdm1^Tg^* mice. **B)** IF detection of CASZ1 (red) and BLIMP1 (green). **C-E**) Quantitation of BLIMP1 (**C**), CASZ1 (**D**), and macromolecular uptake (**E**) in *Villin^CreER^;Prdm1^Tg^* mice. **F)** Cartoon depicting experimental strategy: characterization of neonatal/adult transcription factors and neonatal enterocyte function (macromolecular uptake) in *Ki67^CreER^;Prdm1^Tg^* mice following three days of dex injections. **G-I)** IF analysis of BLIMP1 (green), CASZ1 (red) and macromolecular uptake (magenta) in *Ki67^CreER^;Prdm1^Tg^* mice. Additional image for *Ki67^CreER^;Prdm1^Tg^*mice depicts the same field, but displays macromolecular uptake (Alexa Fluor 488-labeled dextran, magenta) in relation to BLIMP1 expression (green). Quantitation measures CASZ1 levels (**H**) and macromolecular uptake (**I**) in *Prdm1^Tg^-*expressing cells vs. *Prdm1*-negative negative neighboring cells. **J)** Cartoon depicting experimental strategy: characterization of neonatal/adult transcription factors and neonatal enterocyte function (macromolecular uptake) in *Casz1* cKO mice. **K-O)** IF detection (**K**, **L**) and quantitation (**M, N, O**) of CASZ1 (red), BLIMP1 (green), and macromolecular uptake (white). Statistical significance in **H** determined by paired t-test. All other statistical significance determined by unpaired t-test. Scale bars = 25 μm for **G**, 50 μm for **B**, **K**, **L**.

## DISCUSSION

In this study, we identified transcriptional programs that work antagonistically to drive the transition to adulthood. BLIMP1 promotes neonatal enterocyte identity, and maintaining its expression is sufficient to extend neonatal cell function. During maturation, BLIMP1+ neonatal enterocytes are replaced by a wave of adult-like enterocytes in a CASZ1-dependent process. These mutually antagonistic programs act downstream of glucocorticoids: dexamethasone suppresses *Prdm1* and induces *Casz1* expression, and CASZ1 is required for dexamethasone-induced maturation. Together, these findings reveal a dynamic transcriptional switch that controls the transition from suckling to weaning in postnatal intestinal epithelium.

A key finding of this study was the identification of core neonatal and adult gene sets, which reveal conserved metabolic and structural changes underlying the morphological and functional maturation of the intestinal epithelium. The transition between these transcriptional states begins at approximately P14, preceding overt weaning by about a week. This transition proceeds in a single wave from the base of the villus to the tip and is completed just ahead of overt weaning. Maintaining this program until the end of the suckling period may maximize the nutritional benefit provided by milk and support postnatal growth. The timing of this transition is also consistent with prior studies suggesting that maturation is developmentally programmed rather than simply triggered by dietary changes, as delaying weaning has minimal effects on neonatal and adult enzyme expression^13,47^.

Analysis of the core gene sets and further experimentation identified CASZ1 as a heterochronic transcriptional regulator that promotes adult enterocyte fate. Conditional deletion of *Casz1* prolonged neonatal identity and delayed the appearance of adult-like enterocytes, demonstrating that CASZ1 controls the timing of this developmental transition. However, epithelial maturation eventually occurred in *Casz1* cKO mice, indicating that additional factors can compensate for its absence. Thus, CASZ1 is required for the proper timing of intestinal maturation, but is not essential for maturation per se. Heterochronic mutations that alter the timing of developmental transitions have been identified in diverse systems, ranging from plant development to cardiac progenitor differentiation, and can produce lasting defects in tissue morphology and function^48,49^. Whether delayed intestinal maturation similarly has lasting consequences, both within the epithelium and throughout the organism, remains an important question.

Our data further indicate that neonatal and adult enterocyte programs are mutually antagonistic, with CASZ1 and BLIMP1 acting as opposing regulators of cell identity. This is consistent with the morphological and functional differences between neonatal and adult enterocytes, states which cannot co-exist in the same cell. Genetic and pharmacological models that induced maturation promoted CASZ1 expression and eliminated neonatal enterocyte function, including macromolecular uptake. Conversely, maintaining BLIMP1 expression or deleting *Casz1* prolonged neonatal identity. These findings support a model in which the balance between CASZ1 and BLIMP1 governs the transition between neonatal and adult states. This also raises the question of how this switch is initially flipped. The eventual maturation of *Casz1* cKO epithelium further suggests that alternative mechanisms can overcome the neonatal program. Identifying these mechanisms and determining whether the antagonism between CASZ1 and BLIMP1 is direct or mediated through their downstream programs will be important areas for future investigation.

This work identifies glucocorticoids as an upstream hormonal input into the CASZ1/BLIMP1 switch during both spontaneous and dexamethasone-induced maturation. Depletion of epithelial glucocorticoid receptor delayed spontaneous maturation, indicating that endogenous glucocorticoid signaling contributes to the normal timing of this transition. Conversely, dexamethasone suppressed *Prdm1* and induced *Casz1* expression, while either ectopic *Prdm1* expression or conditional deletion of *Casz1* prevented dexamethasone-induced changes in the intestinal epithelium. Thus, glucocorticoids signal directly to the epithelium and act through endogenous transcriptional machinery to promote maturation, providing a mechanistic explanation for observations made more than 40 years ago^14^. However, loss of epithelial glucocorticoid receptor delayed rather than fully prevented maturation, much like deletion of *Casz1*. These findings indicate that glucocorticoids coordinate the timing of the neonatal-to-adult transition through the CASZ1/BLIMP1 switch, while additional inputs can ultimately drive maturation in their absence.

Overall, our data indicate that the neonatal-to-adult transition involves complex mechanisms of mutual antagonism between distinct transcriptional programs. The potential role of hormone signaling on intestinal maturation warrants deeper investigation into the mechanisms that underly neonatal and adult enterocyte identity.

## MATERIALS & METHODS

### Mice

All experiments and procedures were performed under the approval of the Duke Institutional Animal Care and Use Committee (IACUC) and the Duke Laboratory of Animal Research (DLAR). The mouse lines used in this study include CD1 (Charles River), *Villin^Cre/+^* ^50^, *Villin^CreER/+^* (gift from Sylvie Robine, Institut Curie, France), *Prdm1^fl/fl^* ^51^, *MafB^fl/fl^* ^52^; *cMaf^fl/fl^*(aka *Maf^fl/fl^* ^53^), *Casz1^fl/fl^*^41^, *Id3^-/-^* ^36^, *Prdm1^Tg^* (aka *ROSA26:LSL-Prdm1*) ^38^, *Ki67^CreER/+^* ^39^, *Villin^rtTA^* ^45^, *Tre-Degron*^44^, and *GR-GFP*^46^. Both male and female mice were used for all experiments, and genotypes were confirmed by PCR. Mice were maintained in a specific pathogen-free facility, with 12 hour light/dark cycles. For activation of *CreER* alleles, mice were given tamoxifen at a dosage of 50 mg/kg delivered via intraperitoneal (IP) injection (*Villin^CreER^*) or oral gavage (*Ki67^CreER^*). For dexamethasone experiments, mice were given a dose of 0.4 mg/kg delivered via IP injection.

### Isolation of intestinal epithelium

Mice were sacrificed and the small intestine was transferred into cold PBS. The medial region of the small intestine was isolated and flushed with cold PBS to remove luminal contents. The medial small intestine was then opened longitudinally and any remaining luminal contents were removed. Six 1-cm segments of the cleaned small intestine were then transferred to a 5 ml solution of 30 mM EDTA. The exact incubation conditions varied by age: 10 minutes at 37C for P0 – P14 pups, 17 minutes at 37C for P15 – P25 pups, 1-2 hours at 4C rotating for mice older than P25. After the indicated incubation time, the EDTA solution was decanted and tissue pieces were washed briefly and gently with PBS. Tissue pieces were then transferred to a glass petri dish filled with cold PBS, held with forceps, and shaken vigorously to release sheets of epithelial cells from the underlying lamina propria. Epithelial sheets were transferred to a 15 ml conical tube, and processed further according to downstream applications.

### Isolation of single cells from intestinal epithelium

Intestinal epithelial cells were isolated as described above. Isolated epithelial sheets were then added to a 5 ml solution of 0.8 mg/ml dispase, diluted in PBS. All samples were incubated at 37C, and the duration of incubation varied by age: 8 minutes for P0 – P14 mice, 12 min for mice P15 and older. Cells were then mixed by pipetting up and down, and passed through a 70 μm cell strainer into a 50 ml conical tube. Strainers were then rinsed with 10 ml of PBS + 10% FBS. Strained cells were then pelleted by centrifugation, resuspended in PBS + 10% FBS. From this cell resuspension, live cells were quantified and then used for downstream applications.

### RNA isolation and sequencing

A single-cell resuspension of epithelial cells was isolated as described above. Approximately 30 mg of live cells were then spun down for centrifugation, and RNA was isolated by using a Qiagen RNeasy kit (Cat no. 74104) according to the manufacturer’s instructions. RNA was eluted in 35 μl of ultrapure water, and RNA concentration and quality was quantified using a NanoDrop spectrophotometer. Samples were aliquoted and snap frozen, and an aliquot was shipped to Novogene (Sacramento, CA) for RNA sequencing and computational analysis. Unless indicated otherwise in the figure legend, differentially expressed genes were defined by a change in gene expression levels (between control and experimental samples) with an adjusted p-value < 0.05.

### Tissue preparation

Unless otherwise specified, isolated intestinal tissue was either frozen immediately in Optimal Cutting Temperature media (OCT, Sakura) or pre-fixed overnight at 4C in a 4% paraformaldehyde (PFA) solution, diluted in PBS + 0.2% Triton (PBST). For all experiments, proximal, medial, and distal tissue was collected, corresponding to the duodenum, jejunum, and ileum in mature animals. Unless specified otherwise, we focused our analysis on medial tissue for this study. For section analysis, 12 μm tissue sections were collected using a Leica cryostat (CM3050 S). Blocks and additional collected sections were kept at −80C for long-term storage.

### Immunofluorescence

Fresh frozen sections were collected as described above, dried at room temperature (RT), and baked for 10 min at 55C. Sections were then fixed in 4% PFA in PBST for 15 min at room temperature. Fixed sections were then washed three times in PBST, and then blocked for 1 hour in blocking buffer (5% Normal Donkey Serum + 5% Normal Goat Serum + 3% BSA in PBST). Primary antibodies were diluted in blocking buffer (see Supp. Table for dilutions), added to sections, and incubated overnight at 4C. The next day, the primary antibody solution was removed and replaced by a 2 μg/ml Hoescht 34580 stain (diluted in PBST) for 15 min at RT. Slides were then washed three times in PBST, and secondary antibodies diluted in blocking buffer were added to slides for 1 hour at RT. Sections were washed three more times, mounted in mounting media (90% glycerol + 2.5 mg/ml p-phenylenediamine), and sealed with glass coverslips. Immunofluorescent images were acquired using a Zeiss 710 inverted confocal microscope or a Zeiss AxioImager Z1 microscope with an Apotome.2 attachment. Zen software (Zeiss) was used for image acquisition, and image processing was performed using FIJI and photoshop.

### *In vivo* macromolecular uptake assays

To functionally test for neonate-specific macromolecular uptake, neonatal mice were orally gavaged with Alexa Fluor488-conjugated Dextran (ThermoFisher D22910). Mice were sacrificed 3 hours later, and the intestine was isolated and processed as described above. While some endogenous Alexa Fluor488 signal is visible in fresh frozen sections, whenever possible pre-fixed sections of epithelial whole mounts were used for image analysis due to improved clarity of signal.

### Immunostaining of intestinal epithelium wholemounts

The intestinal epithelium was isolated as described above. Sheets of intestinal epithelial cells were transferred to a 1.5 ml tube, and fixed in 4% PFA in PBST for 1 hour, rocking at RT. Samples were then washed three times in PBST, and then resuspended in blocking buffer for 1 hour at RT. After aspirating the blocking buffer, primary antibodies ( diluted in blocking buffer) were added to the samples, and were incubated overnight, rocking, at 4C. The next day, the primary antibody solution was removed and replaced by a 2 μg/ml Hoescht 34580 stain (diluted in PBST) for 15 min at RT. Epithelial cells were then washed three times in PBST, and secondary antibodies diluted in blocking buffer were added to the 1.5 ml tubes for 1 hour at RT. Samples were washed an additional three times with PBST. For mounting, an outline of pre-melted Vaseline, lanolin, and paraffin mix (VALAP) was applied onto a microscopy slide with a paintbrush, and allowed to dry at RT to form a reservoir. Stained epithelial sheets were then transferred into the reservoir, excess PBST was removed, and replaced with mounting media. A coverslip was applied to the top of the reservoir, and was then sealed with an additional layer of VALAP. Immunoflourescence images were then acquired as described previously.

### Edu pulse-chase assays

To label proliferating cells and their progeny, mice were given 5-ethynyl-2’-deoxyuridine (EdU) at a dosage of 2 mg/kg via IP injection. Mice were then sacrificed a set number of hours later, according to the experimental conditions. Injections were staggered to ensure that all animals were collected precisely at the indicated number of hours post-injection. Tissue was then processed for immunofluorescence analysis as described above. After section fixation and subsequent PBST washes, EdU incorporation was visualized using Click-iT chemistry with a Cy5 Picolyl azide (Bioconjugate technologies 1177-1), and imaged using previously-described protocols.

### Quantitation and statistical analysis

For all experiments, a minimum of three biological replicates were used. All animal experiments used a mix of both male and female mice. The statistical tests used to determine significance are indicated in the corresponding figure legend, and were performed in PRISM (Graphpad). For figures displaying quantitative analysis, the following conventions for displaying p-value were used: * p ≤ 0.05, ** p ≤ 0.01, *** p ≤ 0.001, **** p ≤ 0.0001.

## Data and software availability

All sequencing datasets will be made freely available through the NIH Gene Expression Omnibus system (https://www.ncbi.nlm.nih.gov/geo/), and will be released from embargo upon publication.

## ACKNOWLEDGEMENTS

The authors would like to acknowledge and thank the laboratory of Michael Verzi for the generous gift of the *Casz1^fl/fl^* mice. The authors would also like to thank the members of the Lechler and Sumigray laboratories who provided technical assistance and conceptual feedback on this study.

---

**Supplemental Figure 1:**
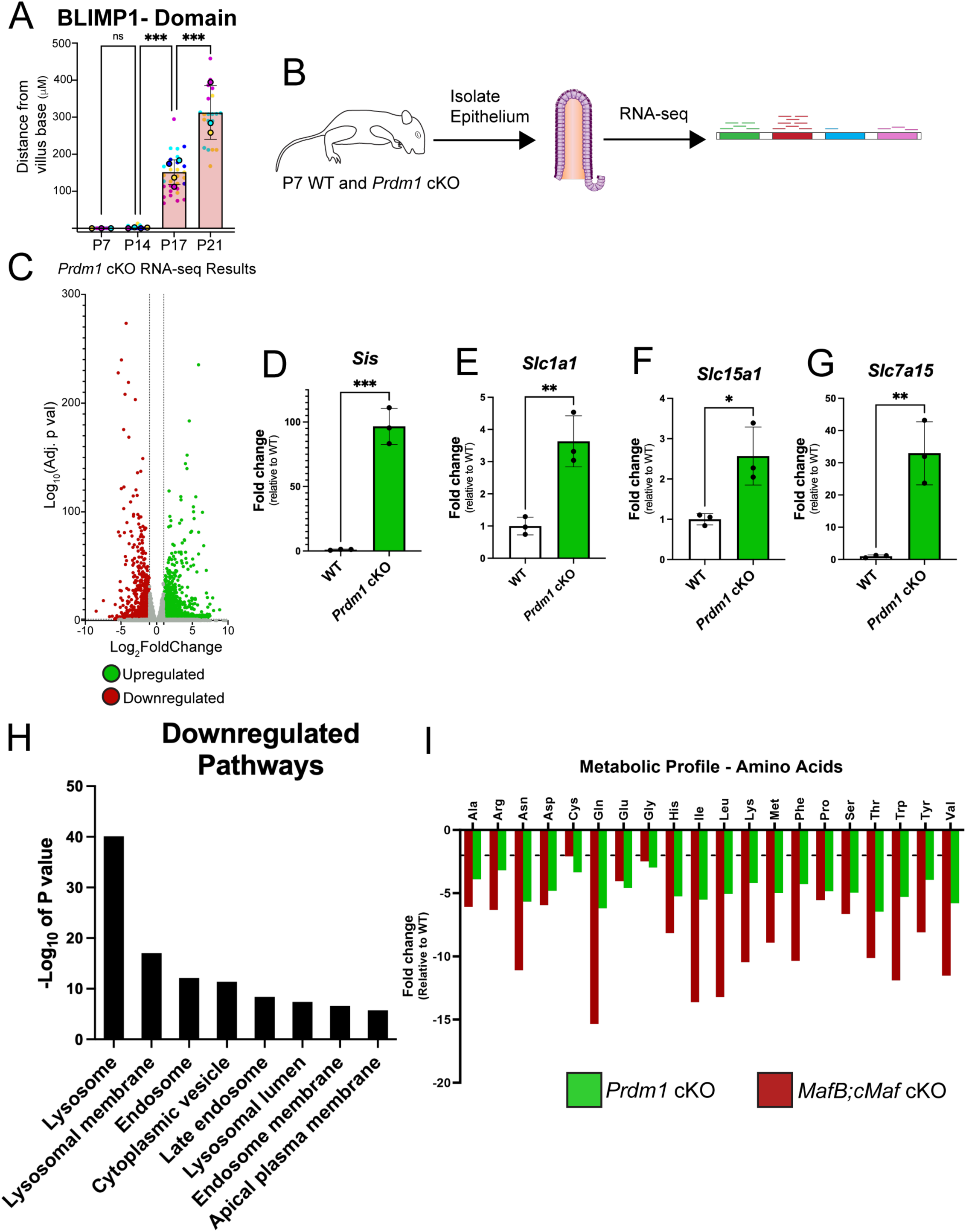
Protein, molecular, and metabolic analysis of WT and *Prdm1* cKO postnatal mice. **A**) Quantitation of the distance from the villus base to BLIMP1-expressing cells (i.e. the length of the BLIMP1-negative domain). Superplot in **A** includes biological replicates (individual mice, circles with border) and the associated technical replicates (individual villi, color-matched open circles). Statistical significance in **A** determined by ordinary one-way ANOVA with Tukey’s multiple comparisons test. **B-G**) RNA-sequencing (RNA-seq) analysis of P7 WT and *Prdm1* cKO neonatal intestinal epithelial cells. Cartoon in **B** depicts experimental strategy. Volcano plot in **C** depicts significantly upregulated (green) and downregulated (red) genes in *Prdm1* cKO neonates. Dotted lines represent p = 0.05 (Y axis) and Log2FoldChange >1 and < −1 (X axis). Bar plots in **D-G** depict gene expression levels of the indicated target genes in WT (white bars) and *Prdm1* cKO (green bars) neonates, as determined by RNA-seq. Gene expression levels determined as the fold change relative to WT (WT mean = 1). Pathway analysis in **H** determined by GO-term analysis. Terms displayed represent a selected set of significantly enriched pathways. **I)** Metabolomic analysis of amino acids in *Prdm1* cKO (green) and *MafB;cMaf* cKO (red) P7 intestinal epithelial cells. Fold change in metabolite concentration is displayed relative to WT littermate controls.

**Supplemental Figure 2:**
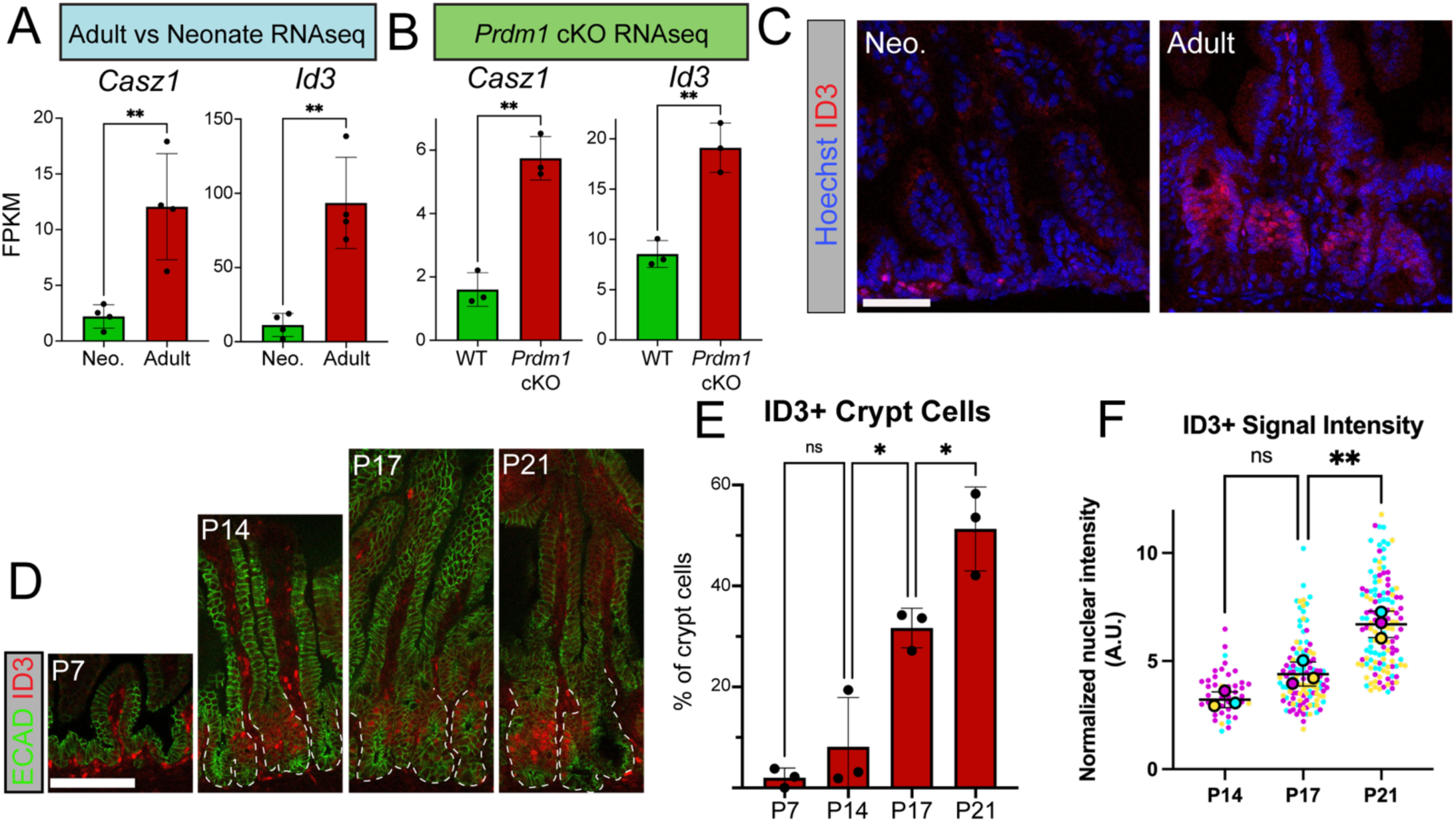
Characterization of *Casz1* and *Id3* expression in postnatal mice. **A-B)** Gene expression levels of epithelial *Casz1* and *Id3* (as determined by RNA-seq) in neonatal vs. adult (**A**, ^1^) and P7 WT vs. *Prdm1* cKO (**B)** mice. Expression levels displayed as FPKM. **C-F)** Characterization of ID3 (red) expression over time. Analysis compares P7 neonates vs. fully mature adults (**C**), as well as detailed time course analysis (**D-F**). Dotted lines in **D** outline crypts. Quantitation (**E-F**) includes percentage of crypt cells that are ID3+ (**E**) and levels of ID3 signal intensity (**F**). Statistical significance determined with ordinary one-way ANOVA with Tukey’s multiple comparisons test. Images in **D** also include IF antibody detection of ECAD (green). **G-H)** Regional IF analysis of CASZ1 expression over time in proximal vs. distal small intestine. Scale bars = 100 μm.

**Supplemental Figure 3:**
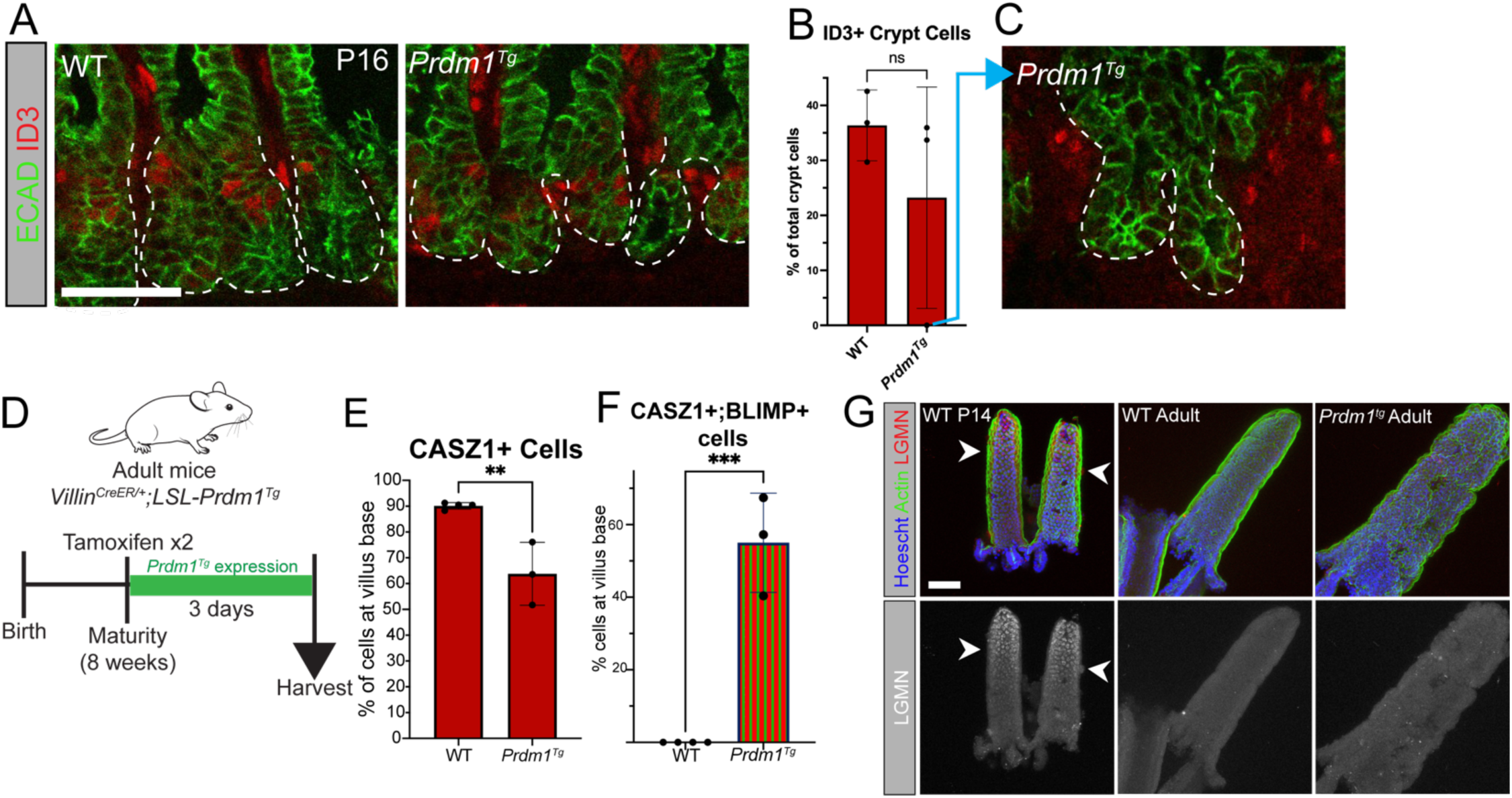
Additional analysis of mice expressing ectopic *Prdm1.* **A-C)** IF antibody detection (**A,C**) and quantitation (**B**) of ID3+ crypt cells in WT and *Prdm1^Tg^* animals. IF images also include ECAD (green). Dotted lines in **A** and **C** outline crypts. Image in **C** demonstrates the lack of epithelial ID3 signal in one of the *Prdm1^Tg^*animals (corresponding quantitation point indicated with blue arrow). **D)** Cartoon depicting experimental strategy for adult activation of *Prdm1^Tg^*. **E)** Quantitation of the percentage of cells that express CASZ1 within 100 μm of villus base. **F)** Percentage of cells (within 100 μm of villus base) that co-express CASZ1/BLIMP1 in *Prdm1^Tg^*mice (or CreER-negative WT littermates). Adult mouse cartoon in **D** acquired from the open source repository SciDraw.io (doi.org/10.5281/zenodo.3925901). **G)** IF analysis of the neonate-specific protease LGMN in neonatal mice (WT P14, positive control), WT adults (negative control), and *Prdm1^Tg^* adults. Arrowheads indicate positive LGMN signal. For all quantitation, statistical significance determined by unpaired t-test. Scale bars = 50 μm.

**Supplemental Figure 4:**
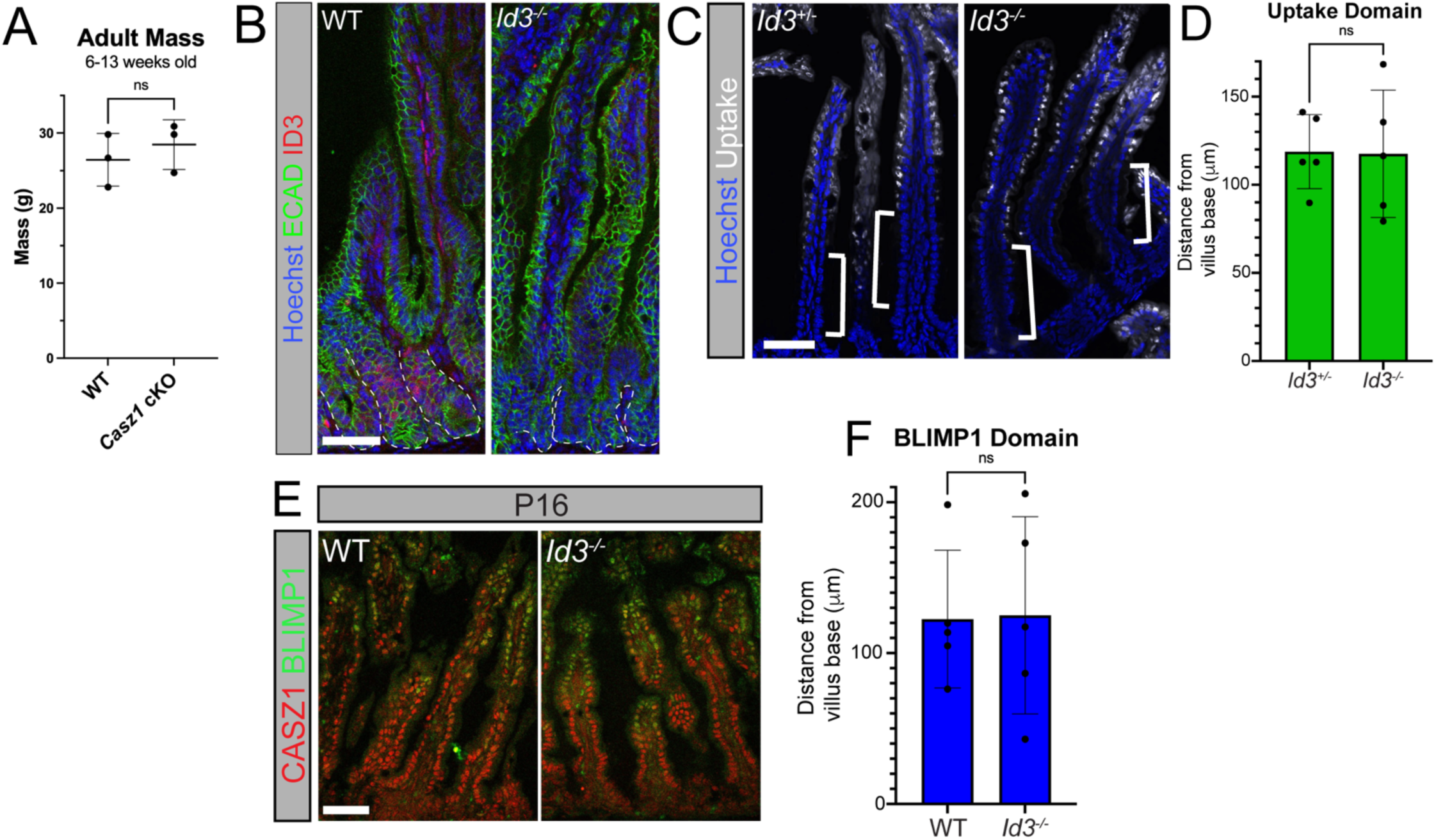
Additional characterization of *Casz1* cKO and *Id3* KO mice. **A)** Mass measurements of *Casz1* cKO adult mice compared to littermate controls. **B)** validation of *Id3^-/-^*mice, as determined by IF antibody detection. Antibody detection includes ID3 (red) and ECAD (green). **C-F**) Functional characterization of P16 Id3^-/-^ mice, as determined by IF. IF detection includes macromolecular uptake (C, Alexa-488-labeled dextran, white), CASZ1 (E, red), and BLIMP1 (E, green). Quantitation in D and F measures the distance from the villus base to the indicated signal. For all quantitation, statistical significance determined by unpaired t-test. Nuclei in B and C marked in blue (Hoescht). Scale bars = 50 μm.

**Supplemental Figure 5:**
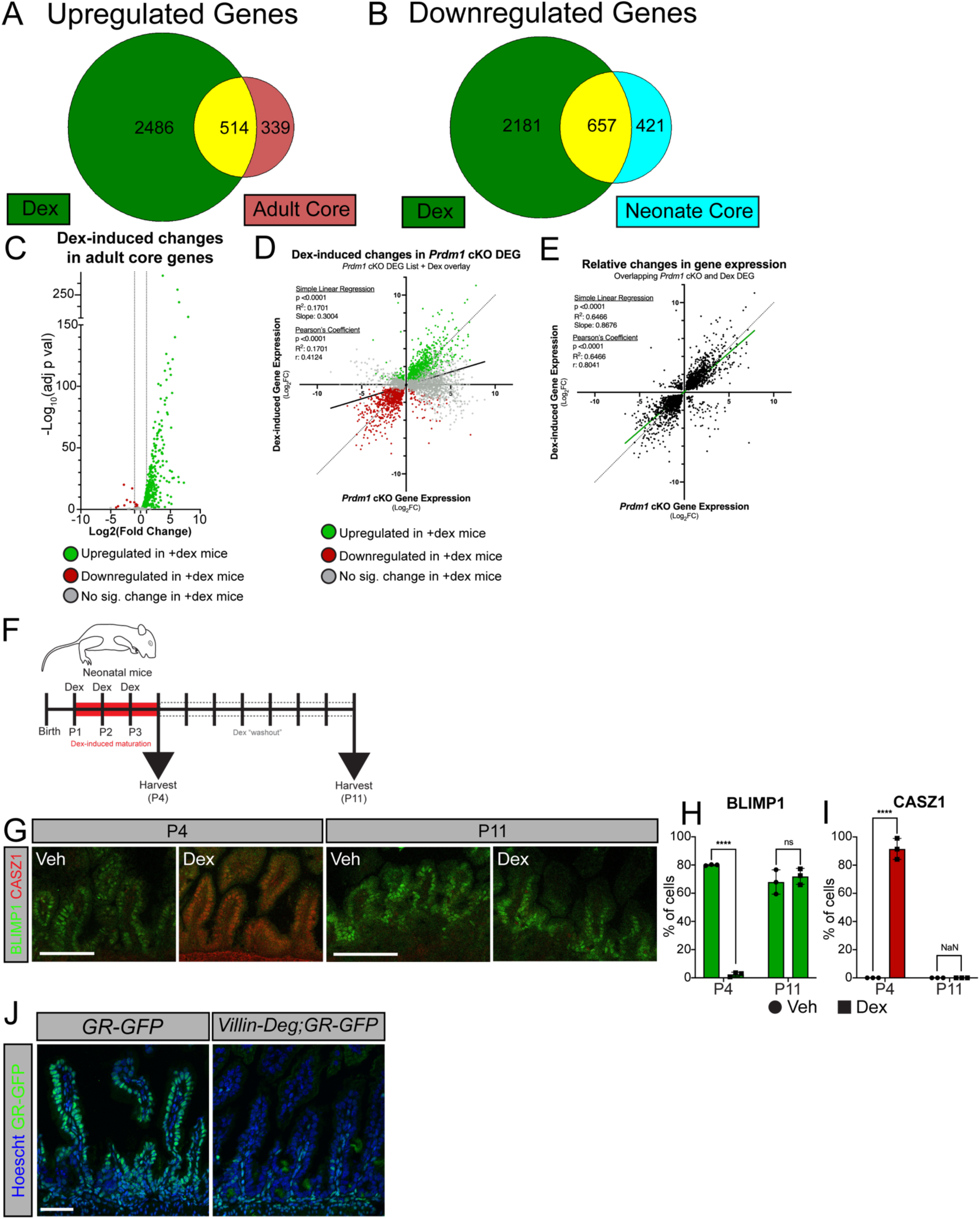
Additional analysis of glucocorticoid-stimulated mice. **A-E)** RNA-sequencing analysis of neonatal (P7) mice following three days (P4, P5, P6) of dex injections. Venn diagrams (**A-B**) compare dex-upregulated genes with the adult core gene list (**A**) and the dex-downregulated list with the neonatal core gene list (**B**). Volcano plot in **C** displays adult core genes that are upregulated in dex-treated animals (green), downregulated (red), or not significantly changed (gray). X-Y analysis in **D** and **E** compares the scale of gene expression changes in dex-injected mice (Y axis) vs. *Prdm1 cKO* mice (X axis). Plot in **D** displays all *Prdm1* cKO differentially expressed genes, and indicates whether these genes were upregulated in dex-treated mice (green), downregulated (red), or had no significant change (gray). Solid black line represents curve of best fit, and the dotted line represents a 1:1 (x = y) linear reference. Plot in **E** only displays overlapping genes between *Prdm1* cKO and Dex-injected mice. **F)** Cartoon depicting experimental strategy: determining if three days of dex injections causes permanent changes in epithelial gene expression. Mice were given three days of dex injection (P1-P3), and were collected at P4 or P11. **G-I)** IF antibody detection (**G**) and quantitation (**H-I**) of BLIMP1 (green) and CASZ1 (red) in Veh/Dex-treated mice at the indicated timepoints. Statistical significance determined by unpaired t-test (timepoints analyzed individually). **J)** Validation of epithelial-specific GR-GFP (green) depletion in *Villin-Deg;GR-GFP*, as determined by IF. Nuclei marked in blue (Hoescht). Scale bars = 50 μm.

## Notes

### Competing Interest Statement

The authors have declared no competing interest.

### Funding

- NIH: R01AR083352, R01AR081081, R35GM150645, F32HD117597, T32DK007568
- Hartwell Foundation: Hartwell Foundation Postdoctoral Research Fellowship

## REFERENCES

1. Hooton, D., Lentle, R., Monro, J., Wickham, M., and Simpson, R. (2015). The Secretion and Action of Brush Border Enzymes in the Mammalian Small Intestine. Rev Physiol Biochem Pharmacol 168, 59–118. 10.1007/112_2015_24.

2. Ferraris, R.P. (2001). Dietary and developmental regulation of intestinal sugar transport. Biochem J 360, 265–276. 10.1042/0264-6021:3600265.

3. Kandasamy, P., Gyimesi, G., Kanai, Y., and Hediger, M.A. (2018). Amino acid transporters revisited: New views in health and disease. Trends Biochem Sci 43, 752–789. 10.1016/j.tibs.2018.05.003.

4. Skrzypek, T., Valverde Piedra, J.L., Skrzypek, H., Kazimierczak, W., Biernat, M., and Zabielski, R. (2007). Gradual disappearance of vacuolated enterocytes in the small intestine of neonatal piglets. J Physiol Pharmacol 58 Suppl 3, 87–95.

5. Clark, S.L., Jr. (1959). The ingestion of proteins and colloidal materials by columnar absorptive cells of the small intestine in suckling rats and mice. J Biophys Biochem Cytol 5, 41–50. 10.1083/jcb.5.1.41.

6. Cornell, R., and Padykula, H.A. (1969). A cytological study of intestinal absorption in the suckling rat. Am J Anat 125, 291–315. 10.1002/aja.1001250304.

7. Bara, A.M., Chen, L., Ma, C., Underwood, J., Moreci, R.S., Sumigray, K., Sun, T., Diao, Y., Verzi, M., and Lechler, T. (2022). Maf family transcription factors are required for nutrient uptake in the mouse neonatal gut. Development 149. 10.1242/dev.201251.

8. Park, J., Levic, D.S., Sumigray, K.D., Bagwell, J., Eroglu, O., Block, C.L., Eroglu, C., Barry, R., Lickwar, C.R., Rawls, J.F., et al. (2019). Lysosome-Rich Enterocytes Mediate Protein Absorption in the Vertebrate Gut. Dev Cell 51, 7–20 e26. 10.1016/j.devcel.2019.08.001.

9. Wilson, J.M., Whitney, J.A., and Neutra, M.R. (1991). Biogenesis of the apical endosome-lysosome complex during differentiation of absorptive epithelial cells in rat ileum. J Cell Sci 100 (Pt 1), 133–143. 10.1242/jcs.100.1.133.

10. Wissig, S.L., and Graney, D.O. (1968). Membrane modifications in the apical endocytic complex of ileal epithelial cells. J Cell Biol 39, 564–579. 10.1083/jcb.39.3.564.

11. Rath, T., Baker, K., Pyzik, M., and Blumberg, R.S. (2014). Regulation of immune responses by the neonatal fc receptor and its therapeutic implications. Front Immunol 5, 664. 10.3389/fimmu.2014.00664.

12. Gericke, B., Amiri, M., and Naim, H.Y. (2016). The multiple roles of sucrase-isomaltase in the intestinal physiology. Mol Cell Pediatr 3, 2. 10.1186/s40348-016-0033-y.

13. Henning, S.J. (1981). Postnatal development: coordination of feeding, digestion, and metabolism. Am J Physiol 241, G199–214. 10.1152/ajpgi.1981.241.3.G199.

14. Henning, S.J. (1985). Ontogeny of enzymes in the small intestine. Annu Rev Physiol 47, 231–245. 10.1146/annurev.ph.47.030185.001311.

15. Harper, J., Mould, A., Andrews, R.M., Bikoff, E.K., and Robertson, E.J. (2011). The transcriptional repressor Blimp1/Prdm1 regulates postnatal reprogramming of intestinal enterocytes. Proc Natl Acad Sci U S A 108, 10585–10590. 10.1073/pnas.1105852108.

16. Muncan, V., Heijmans, J., Krasinski, S.D., Buller, N.V., Wildenberg, M.E., Meisner, S., Radonjic, M., Stapleton, K.A., Lamers, W.H., Biemond, I., et al. (2011). Blimp1 regulates the transition of neonatal to adult intestinal epithelium. Nat Commun 2, 452. 10.1038/ncomms1463.

17. Solomon, N.S., Gartner, H., Oesterreicher, T.J., and Henning, S.J. (2001). Development of glucocorticoid-responsiveness in mouse intestine. Pediatr Res 49, 782–788. 10.1203/00006450-200106000-00012.

18. Martin, G.R., and Henning, S.J. (1982). Relative importance of corticosterone and thyroxine in the postnatal development of sucrase and maltase in rat small intestine. Endocrinology 111, 912–918. 10.1210/endo-111-3-912.

19. Henning, S.J., and Sims, J.M. (1979). Delineation of the glucocorticoid-sensitive period of intestinal development in the rat. Endocrinology 104, 1158–1163. 10.1210/endo-104-4-1158.

20. Nanthakumar, N.N., and Henning, S.J. (1993). Ontogeny of sucrase-isomaltase gene expression in rat intestine: responsiveness to glucocorticoids. Am J Physiol 264, G306–311. 10.1152/ajpgi.1993.264.2.G306.

21. Herbst, J.J., and Koldovsky, O. (1972). Cell migration and cortisone induction of sucrase activity in jejunum and ileum. Biochem J 126, 471–476. 10.1042/bj1260471.

22. Koldovsky, O., Jirsova, V., and Heringova, A. (1965). Effect of aldosterone and corticosterone on beta-galactosidase and invertase activity in the small intestine of rats. Nature 206, 300–301. 10.1038/206300a0.

23. Koldovsky, O., Jumawan, J., and Palmieri, M. (1975). Effect of thyroidectomy on the activity of alpha-glucosidases and acid hydrolases in the small intestine of rats during weaning. J Endocrinol 66, 31–36. 10.1677/joe.0.0660031.

24. Yeh, K.Y., and Moog, M. (1977). Influence of the thyroid and adrenal glands on the growth of the intestine of the suckling rat, and on the development of intestinal alkaline phosphatase and disaccharidase activities. J Exp Zool 200, 337–347. 10.1002/jez.1402000304.

25. Committee on Obstetric, P. (2017). Committee Opinion No. 713: Antenatal Corticosteroid Therapy for Fetal Maturation. Obstet Gynecol 130, e102–e109. 10.1097/AOG.0000000000002237.

26. Cogo, P.E., Simonato, M., Mariatoffolo, G., Stefanutti, G., Chierici, M., Cobelli, C., Ori, C., and Carnielli, V.P. (2008). Dexamethasone therapy in preterm infants developing bronchopulmonary dysplasia: effect on pulmonary surfactant disaturated-phosphatidylcholine kinetics. Pediatr Res 63, 433–437. 10.1203/PDR.0b013e3181659759.

27. Navis, M., Martins Garcia, T., Renes, I.B., Vermeulen, J.L., Meisner, S., Wildenberg, M.E., van den Brink, G.R., van Elburg, R.M., and Muncan, V. (2019). Mouse fetal intestinal organoids: new model to study epithelial maturation from suckling to weaning. EMBO Rep 20. 10.15252/embr.201846221.

28. van der Flier, L.G., and Clevers, H. (2009). Stem cells, self-renewal, and differentiation in the intestinal epithelium. Annu Rev Physiol 71, 241–260. 10.1146/annurev.physiol.010908.163145.

29. Gehart, H., and Clevers, H. (2019). Tales from the crypt: new insights into intestinal stem cells. Nat Rev Gastroenterol Hepatol 16, 19–34. 10.1038/s41575-018-0081-y.

30. Kodama, N., Iwao, T., Katano, T., Ohta, K., Yuasa, H., and Matsunaga, T. (2016). Characteristic Analysis of Intestinal Transport in Enterocyte-Like Cells Differentiated from Human Induced Pluripotent Stem Cells. Drug Metab Dispos 44, 0. 10.1124/dmd.116.069336.

31. Hu, Q.X., Ottestad-Hansen, S., Holmseth, S., Hassel, B., Danbolt, N.C., and Zhou, Y. (2018). Expression of Glutamate Transporters in Mouse Liver, Kidney, and Intestine. J Histochem Cytochem 66, 189–202. 10.1369/0022155417749828.

32. Cosovanu, C., Resch, P., Jordan, S., Lehmann, A., Ralser, M., Farztdinov, V., Spranger, J., Mulleder, M., Brachs, S., and Neumann, C. (2022). Intestinal epithelial c-Maf expression determines enterocyte differentiation and nutrient uptake in mice. J Exp Med 219. 10.1084/jem.20220233.

33. Zwick, R.K., Kasparek, P., Palikuqi, B., Viragova, S., Weichselbaum, L., McGinnis, C.S., McKinley, K.L., Rathnayake, A., Vaka, D., Nguyen, V., et al. (2024). Epithelial zonation along the mouse and human small intestine defines five discrete metabolic domains. Nat Cell Biol 26, 250–262. 10.1038/s41556-023-01337-z.

34. Block, C.L., Childers, L., Cortez, A.L., Sakers, K., Lewis, T.R., Levic, D.S., Frazer, L.C., Jania, C.M., Arshavsky, V.Y., Good, M., et al. (2025). The mouse neonatal small intestine is regionally specialized for protein absorption and transepithelial transport. Development 152. 10.1242/dev.205127.

35. Mattar, P., Ericson, J., Blackshaw, S., and Cayouette, M. (2015). A conserved regulatory logic controls temporal identity in mouse neural progenitors. Neuron 85, 497–504. 10.1016/j.neuron.2014.12.052.

36. Pan, L., Sato, S., Frederick, J.P., Sun, X.H., and Zhuang, Y. (1999). Impaired immune responses and B-cell proliferation in mice lacking the Id3 gene. Mol Cell Biol 19, 5969–5980. 10.1128/MCB.19.9.5969.

37. Yang, C.Y., Best, J.A., Knell, J., Yang, E., Sheridan, A.D., Jesionek, A.K., Li, H.S., Rivera, R.R., Lind, K.C., D’Cruz, L.M., et al. (2011). The transcriptional regulators Id2 and Id3 control the formation of distinct memory CD8+ T cell subsets. Nat Immunol 12, 1221–1229. 10.1038/ni.2158.

38. Goodson, N.B., Park, K.U., Silver, J.S., Chiodo, V.A., Hauswirth, W.W., and Brzezinski, J.A.t. (2020). Prdm1 overexpression causes a photoreceptor fate-shift in nascent, but not mature, bipolar cells. Dev Biol 464, 111–123. 10.1016/j.ydbio.2020.06.003.

39. Basak, O., Krieger, T.G., Muraro, M.J., Wiebrands, K., Stange, D.E., Frias-Aldeguer, J., Rivron, N.C., van de Wetering, M., van Es, J.H., van Oudenaarden, A., et al. (2018). Troy+ brain stem cells cycle through quiescence and regulate their number by sensing niche occupancy. Proc Natl Acad Sci U S A 115, E610–E619. 10.1073/pnas.1715911114.

40. Malagola, E., Vasciaveo, A., Ochiai, Y., Kim, W., Zheng, B., Zanella, L., Wang, A.L.E., Middelhoff, M., Nienhuser, H., Deng, L., et al. (2024). Isthmus progenitor cells contribute to homeostatic cellular turnover and support regeneration following intestinal injury. Cell 187, 3056–3071 e3017. 10.1016/j.cell.2024.05.004.

41. Dorr, K.M., Amin, N.M., Kuchenbrod, L.M., Labiner, H., Charpentier, M.S., Pevny, L.H., Wessels, A., and Conlon, F.L. (2015). Casz1 is required for cardiomyocyte G1-to-S phase progression during mammalian cardiac development. Development 142, 2037–2047. 10.1242/dev.119107.

42. Nanthakumar, N.N., and Henning, S.J. (1995). Distinguishing normal and glucocorticoid-induced maturation of intestine using bromodeoxyuridine. Am J Physiol 268, G139–145. 10.1152/ajpgi.1995.268.1.G139.

43. Agbemafle, B.M., Oesterreicher, T.J., Shaw, C.A., and Henning, S.J. (2005). Immediate early genes of glucocorticoid action on the developing intestine. Am J Physiol Gastrointest Liver Physiol 288, G897–906. 10.1152/ajpgi.00454.2004.

44. Prado-Mantilla, A., Sheheen, J., Underwood, J., and Lechler, T. (2026). Cell type-specific spatiotemporal control of GFP-tagged protein degradation in mice. J Cell Biol 225. 10.1083/jcb.202504003.

45. Chen, Y.J., Finkbeiner, S.R., Weinblatt, D., Emmett, M.J., Tameire, F., Yousefi, M., Yang, C., Maehr, R., Zhou, Q., Shemer, R., et al. (2014). De novo formation of insulin-producing "neo-beta cell islets" from intestinal crypts. Cell Rep 6, 1046–1058. 10.1016/j.celrep.2014.02.013.

46. Brewer, J.A., Sleckman, B.P., Swat, W., and Muglia, L.J. (2002). Green fluorescent protein-glucocorticoid receptor knockin mice reveal dynamic receptor modulation during thymocyte development. J Immunol 169, 1309–1318. 10.4049/jimmunol.169.3.1309.

47. Motohashi, Y., Fukushima, A., Kondo, T., and Sakuma, K. (1997). Lactase decline in weaning rats is regulated at the transcriptional level and not caused by termination of milk ingestion. J Nutr 127, 1737–1743. 10.1093/jn/127.9.1737.

48. Lee, D.Y., Lee, J., Moon, S., Park, S.Y., and An, G. (2007). The rice heterochronic gene SUPERNUMERARY BRACT regulates the transition from spikelet meristem to floral meristem. Plant J 49, 64–78. 10.1111/j.1365-313X.2006.02941.x.

49. Rowton, M., Perez-Cervantes, C., Hur, S., Jacobs-Li, J., Lu, E., Deng, N., Guzzetta, A., Hoffmann, A.D., Stocker, M., Steimle, J.D., et al. (2022). Hedgehog signaling activates a mammalian heterochronic gene regulatory network controlling differentiation timing across lineages. Dev Cell 57, 2181–2203 e2189. 10.1016/j.devcel.2022.08.009.

50. Madison, B.B., Dunbar, L., Qiao, X.T., Braunstein, K., Braunstein, E., and Gumucio, D.L. (2002). Cis elements of the villin gene control expression in restricted domains of the vertical (crypt) and horizontal (duodenum, cecum) axes of the intestine. J Biol Chem 277, 33275–33283. 10.1074/jbc.M204935200.

51. Shapiro-Shelef, M., Lin, K.I., McHeyzer-Williams, L.J., Liao, J., McHeyzer-Williams, M.G., and Calame, K. (2003). Blimp-1 is required for the formation of immunoglobulin secreting plasma cells and pre-plasma memory B cells. Immunity 19, 607–620. 10.1016/s1074-7613(03)00267-x.

52. Yu, W.M., Appler, J.M., Kim, Y.H., Nishitani, A.M., Holt, J.R., and Goodrich, L.V. (2013). A Gata3-Mafb transcriptional network directs post-synaptic differentiation in synapses specialized for hearing. Elife 2, e01341. 10.7554/eLife.01341.

53. Wende, H., Lechner, S.G., Cheret, C., Bourane, S., Kolanczyk, M.E., Pattyn, A., Reuter, K., Munier, F.L., Carroll, P., Lewin, G.R., and Birchmeier, C. (2012). The transcription factor c-Maf controls touch receptor development and function. Science 335, 1373–1376. 10.1126/science.1214314.

